# Schlafen family member 5 (SLFN5) regulates LAT1-mediated mTOR activation in castration-resistant prostate cancer

**DOI:** 10.1101/2020.09.17.301283

**Authors:** Rafael S. Martinez, Mark J. Salji, Linda Rushworth, Chara Ntala, Giovanny Rodriguez Blanco, Ann Hedley, William Clark, Paul Peixoto, Eric Hervouet, Elodie Renaude, Sonia H.Y. Kung, Laura C.A. Galbraith, Colin Nixon, Sergio Lilla, Gillian M. MacKay, Ladan Fazli, David Sumpton, Martin E. Gleave, Sara Zanivan, Arnaud Blomme, Hing Y. Leung

**Affiliations:** CRUK Beatson Institute, Garscube Estate, Switchback Road, Glasgow, G61 1BD, UK; Institute of Cancer Sciences, University of Glasgow, Garscube Estate, Switchback Road, Glasgow, G61 1QH, UK; Univ. Bourgogne Franche-Comté, INSERM, EFS BFC, UMR1098, Interactions Hôte-Greffon-Tumeur/Ingénierie Cellulaire et Génique, F-25000 Besançon, France; EPIGENExp, (EPIgenetics and GENe EXPression Technical Platform), Besançon France; Department of Urologic Sciences, Faculty of Medicine, University of British Columbia, Vancouver, BC V5Z 1M9, Canada; Vancouver Prostate Centre, Vancouver, BC V6H 3Z6, Canada

**Keywords:** castration-resistant prostate cancer, proteomics, cancer metabolism, LAT1, mTOR signalling

## Abstract

Androgen-deprivation therapy (ADT) is the standard of care for the treatment of non-resectable prostate cancer (PCa). Despite high treatment efficiency, most patients ultimately develop lethal castration-resistant prostate cancer (CRPC). In this study, we perform a comparative proteomic analysis of three *in vivo,* androgen receptor (AR)–driven, orthograft models of CRPC. Differential proteomic analysis reveals that distinct molecular mechanisms, including amino acid (AA) and fatty acid (FA) metabolism, are involved in the response to ADT between the different models. Despite this heterogeneity, we identify SLFN5 as an AR-regulated biomarker in CRPC. SLFN5 expression is high in CRPC tumours and correlates with poor patient outcome. *In vivo, SLFN5* depletion strongly impairs tumour growth in castrated condition. Mechanistically, SLFN5 interacts with ATF4 and regulates the expression of LAT1, an essential AA transporter. Consequently, *SLFN5* depletion in CRPC cells decreases intracellular levels of essential AA and impairs mTORC1 signalling in a LAT1-dependent manner.

## Introduction

Androgen deprivation therapy (ADT), together with direct targeting of the androgen receptor (AR) pathway, remains the most effective treatment for patients with advanced prostate cancer (PCa). However, patients that relapse will ultimately develop a lethal form of the disease, termed castration-resistant prostate cancer (CRPC), with current second line therapeutic options providing only relatively short gain in survival. Therefore, a better understanding of the molecular mechanisms underlying treatment resistance and the identification of specific CRPC markers remain a subject of intensive research focus.

Targeting cancer metabolism, using small molecule inhibitors or diet manipulation, alone or in combination with existing drugs, represents an appealing option to further refine anti-cancer therapies^1^. Due to the basal metabolism of the prostate gland, PCa is associated with distinct metabolic features, such as a reliance on oxidative phosphorylation^2^ in the early stage of the disease. Progression to CRPC, as well as resistance to treatment, is often accompanied with a metabolic switch that renders PCa tumours increasingly dependent on specific metabolic pathways such as glycolysis^3^, lipid and cholesterol metabolism^4^. Alteration of cancer cell metabolism can result from the activation of multiple signalling pathways, which are often strongly inter-connected between each other. In prostate, AR has been shown to directly control glucose and lipid metabolism of cancer cells, thus supporting cancer progression^5^ or treatment resistance^6^. In addition to AR, mTORC1 signalling is frequently dysregulated in PCa^7^. Regulation of cellular metabolism and protein synthesis by mTORC1 is critical to sustain the biomass required for enhanced proliferation in cancer cells. However, the limited success achieved by current mTOR inhibitors in clinics points towards the need to better characterise other factors upstream of mTOR regulation^8^.

Along with growth factors, amino acid (AA) homeostasis is essential for the regulation of mTORC1 activity. Leucine, in particular, is critical for mTORC1 localisation at the surface of lysosomes^9^. Thus, an important component of this metabolic process is the L-type amino acid transporter LAT1. Mechanistically, LAT1 mediates the intracellular uptake of branched chain and aromatic AA in exchange for glutamine, in a sodium-independent manner^10^. LAT1 over-expression has been reported in multiple cancer types, including PCa^11^. In patients, LAT1 expression is elevated following ADT and in metastatic lesions^12,13^. Mechanistically, LAT1 is regulated by the stress-induced transcription factor ATF4 and contributes to PCa progression, at least in part, by sustaining mTORC1 signalling^13^.

Schlafen family member 5 (SLFN5) is a member of the Schlafen family of proteins, a group of type 1 interferon-inducible proteins. In addition to an AAA (ATPase) domain and a specific SLFN box, the *SLFN5* gene contains a helicase domain as well as a nuclear localisation sequence, which suggests a role for this protein in transcription-related processes^14^. However, the molecular function of SLFN5, as well as its contribution to cancer, remains unclear. SLFN5 levels correlate with good patient outcome in melanoma^15^, breast^16^ and renal cancers^17^, and SLFN5 expression was associated with decreased cell motility in these cell types. A recent study further reported that SLFN5 negatively regulates invasiveness and epithelial-to-mesenchymal transition in breast cancer cells by directly controlling the transcription of ZEB1^18^. By contrast, a pro-tumourigenic role for SLFN5 has been suggested in glioblastoma, where SLFN5 acts as a transcriptional co-repressor of STAT1 following type 1-interferon treatment^19^. Taken together, these results suggest that the role of SLFN5 in cancer progression might be context-dependent.

In this study, we develop and characterise three *in vivo*, AR-driven, orthograft models of PCa that accurately model patient CRPC condition. In depth proteomic analysis reveals a complex response to hormone deprivation therapy, indicating distinct molecular mechanisms across the different models. Despite this molecular heterogeneity, we identify SLFN5 as a common target, the expression of which is consistently up-regulated upon ADT resistance. SLFN5 expression is increased in treatment-resistant patient biopsies, while SLFN5 deletion dramatically impairs the growth of CRPC tumours *in vivo*. Mechanistically, we show that SLFN5 directly interacts with ATF4 and strongly controls the expression of several ATF4-enriched target genes, including the AA transporter LAT1. Consequently, we demonstrate that SLFN5 knockout (KO) in CRPC cells alters AA metabolism and disrupts mTORC1 signalling in a LAT1-dependent manner, presenting a potential therapeutic target.

## Results

### Proteomic characterisation of *in vivo* models of CRPC

To study resistance to androgen deprivation, we developed three independent PCa orthograft models to mimic clinical CRPC by injecting matched pairs (hormone naïve and castration resistant) of AR-proficient human PCa cell lines into the prostate of immuno-deficient mice. In CRPC conditions, orthotopic injection was directly followed by orchidectomy to achieve ADT. LNCaP, CWR22res and VCaP were selected based on their differences in AR expression, full length and variants, and herein referred to as hormone naïve cells (HN) (Supplementary figure 1a). HN cells were cultured *in vitro* in androgen-containing medium (supplemented with foetal bovine serum) and injected orthotopically into the prostate of uncastrated mice. By contrast, two matched, isogenic, androgen-independent (CRPC) cell lines, namely LNCaP AI and 22rv1, were routinely cultured *in vitro* in androgen-deficient medium (supplemented with charcoal-stripped serum) and orthotopically injected into castrated mice. VCaP cells, which were able to grow orthotopically in castrated mice, were injected into both uncastrated and castrated mice (Figure 1a). All models develop CRPC tumours *in vivo* and have been individually used in the literature^20^. Surprisingly, the castrate models of LNCaP AI and VCaP-CR orthografts displayed a higher incidence than their HN counterparts. By contrast, CWR22res were more tumourigenic than 22rv1 (Supplementary Figure 1b). Importantly, there was no significant difference in the final tumour weight between HN and CRPC models (Supplementary Figure 1c).

**Figure 1:**
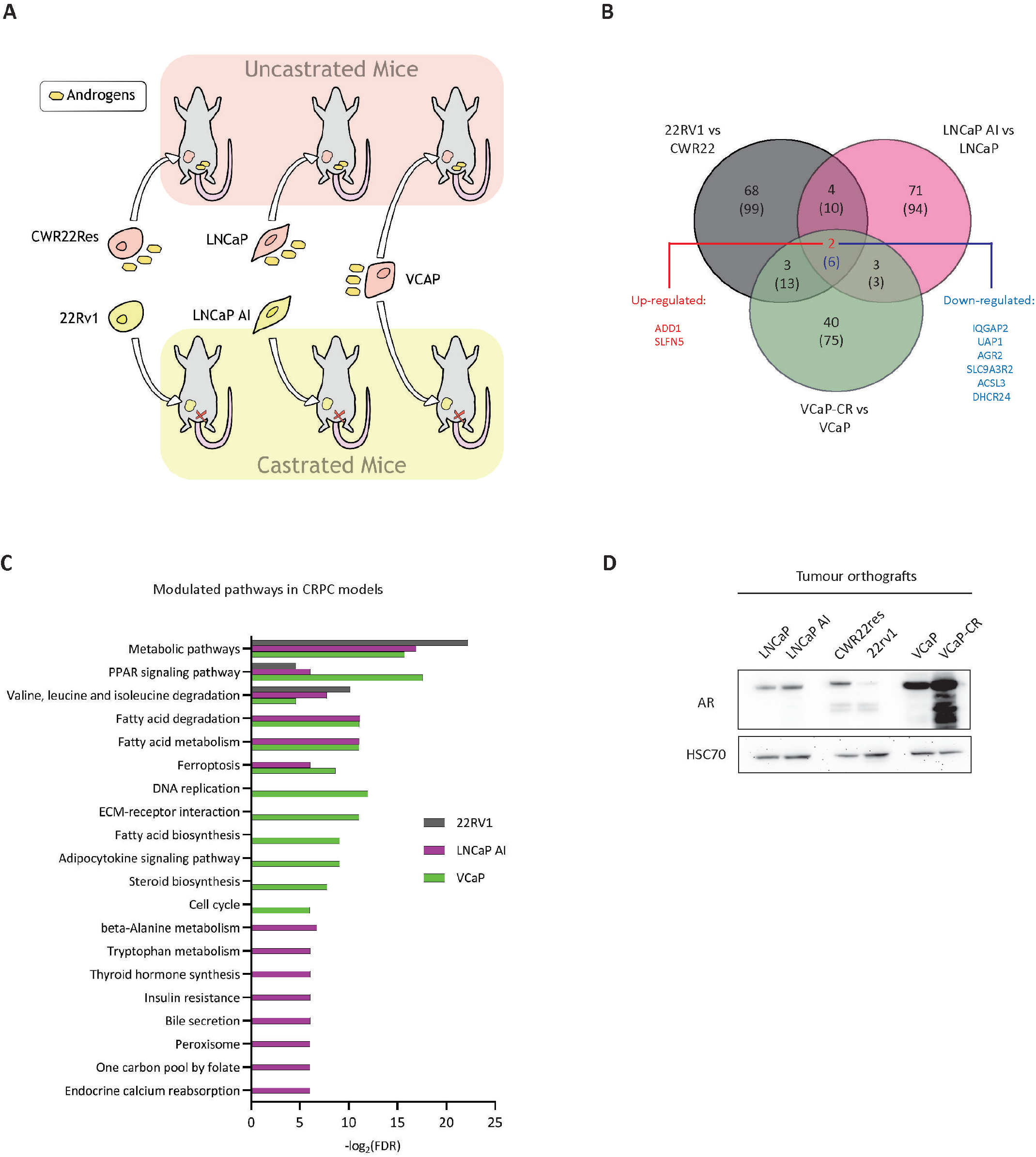
Proteomic characterisation of *in vivo* CRPC orthografts. **A**, Schematic representation of the three CRPC orthograft models used in this study. **B**, Venn diagrams highlighting proteins commonly modulated (p-value < 0.05, FC = 2) in CRPC orthografts in comparison to their respective HN counterparts. Up-regulated proteins are on top; Down-regulated proteins are into brackets. **C**, Top 20 enriched pathways (KEGG pathways) significantly modulated in the proteomic analysis of CRPC orthografts. Pathway enrichment analysis was performed using the STRING database (http://string-db.org). **D**, Western blot analysis of AR expression in HN and matched CRPC tumour orthografts. HSC70 was used as a sample loading control.

Using a SILAC-based proteomic approach, we compared the proteomes of the CRPC and HN tumours within each model. This allowed us to define three independent proteomic signatures associated with CRPC (Figure 1b, Supplementary Data 1). Strikingly, enrichment pathway analysis highlighted changes in metabolism as the top pathway commonly modulated in CRPC (Figure 1c). In particular, pathways related to lipid (PPAR signalling, fatty acid oxidation, ferroptosis) and amino acid metabolism (branched chain AA degradation) were significantly modulated upon ADT resistance (Figure 1c). However, the regulated proteins involved in these pathways varied greatly among the models, emphasising the molecular heterogeneity of CRPC (Supplementary Data 1). For instance, the 22rv1 model was characterised by enrichment in EGFR signalling. This EGFR signature was also observed to a lesser extent in the LNCaP AI model, but not in the VCaP-CR. LNCaP AI tumours were characterised by an increased expression of several components of the mitochondrial electron transport chain as well as of the unfolded protein response, while the VCaP-CR tumours displayed down-regulation of a large cluster of mitosis-associated proteins (Supplementary Data 1). This heterogeneity was further exemplified by different patterns of AR expression following ADT across the different models (decreased in 22rv1 when compared to CWR22res, increased in LNCaP AI compared to LNCaP, and strongly increased in VCaP-CR tumours in comparison to VCAP, Figure 1d). Finally, we took advantage of our proteomic approach to generate a proteomic signature characteristic of CRPC, irrespective of tumour type or AR status. We compared our proteomic datasets to identify proteins that were commonly modulated in all three models of CRPC. Interestingly, only 8 proteins were commonly regulated across the different models (FC = 2; p-value < 0.05; Figure 1b). Among these candidates, Adducin-1 (ADD1) and Schlafen Family Member 5 (SLFN5) were significantly more abundant in all CRPC tumours. Unlike ADD1, the role of SLFN5 in cancer remains understudied, which prompted us to explore its function in CRPC.

### *SLFN5* is an AR-regulated gene highly expressed in CRPC

We first confirmed high SLFN5 levels in CRPC orthografts by performing western blot on total tumour lysates (Figure 2a). Immunohistochemistry performed on tumour slides further evidenced a strong nuclear staining for SLFN5 in epithelial cells. In agreement with the proteomic data, SLFN5 staining was more intense in 22rv1 and LNCaP AI tumours, when compared to their HN counterparts (Figure 2b). Increased SLFN5 expression was also observed *in vitro* following long-term androgen deprivation (Figure 2c). AR is the main driver of ADT resistance^6^ and directly regulates the expression of multiple genes involved in a plethora of biological processes that, if aberrantly regulated, are known to cause cancer pathogenesis^21^. Therefore we tested the ability of AR to modulate SLFN5 expression in PCa cells. In LNCaP, short-term androgen deprivation (72 h) was sufficient to increase *SLFN5* mRNA by almost four-fold, and this effect was partially rescued by the addition of dihydrotestosterone (DHT) (Figure 2d). In CRPC cells, short-term addition of DHT also decreased SLFN5 level in a dose-dependent manner (Figure 2e). In general, SLFN5 expression was increased upon androgen withdrawal and showed an inverse correlation with AR expression (Figure 2f). Transient silencing of AR using siRNA increased *SLFN5* mRNA expression in both LNCaP and CWR22res cells (Figure 2g). Interestingly, we also observed that *SLFN5* expression was inversely correlated to the expression of the canonical AR target gene *KLK3* in prostate cancer patients (using the TCGA PRAD dataset, Figure 2h). Finally, we confirmed the binding of AR on the promoter region of *SLFN5* using chromatin immunoprecipitation (ChIP) (Figure 2i), thus validating SLFN5 as a direct AR target in PCa.

**Figure 2:**
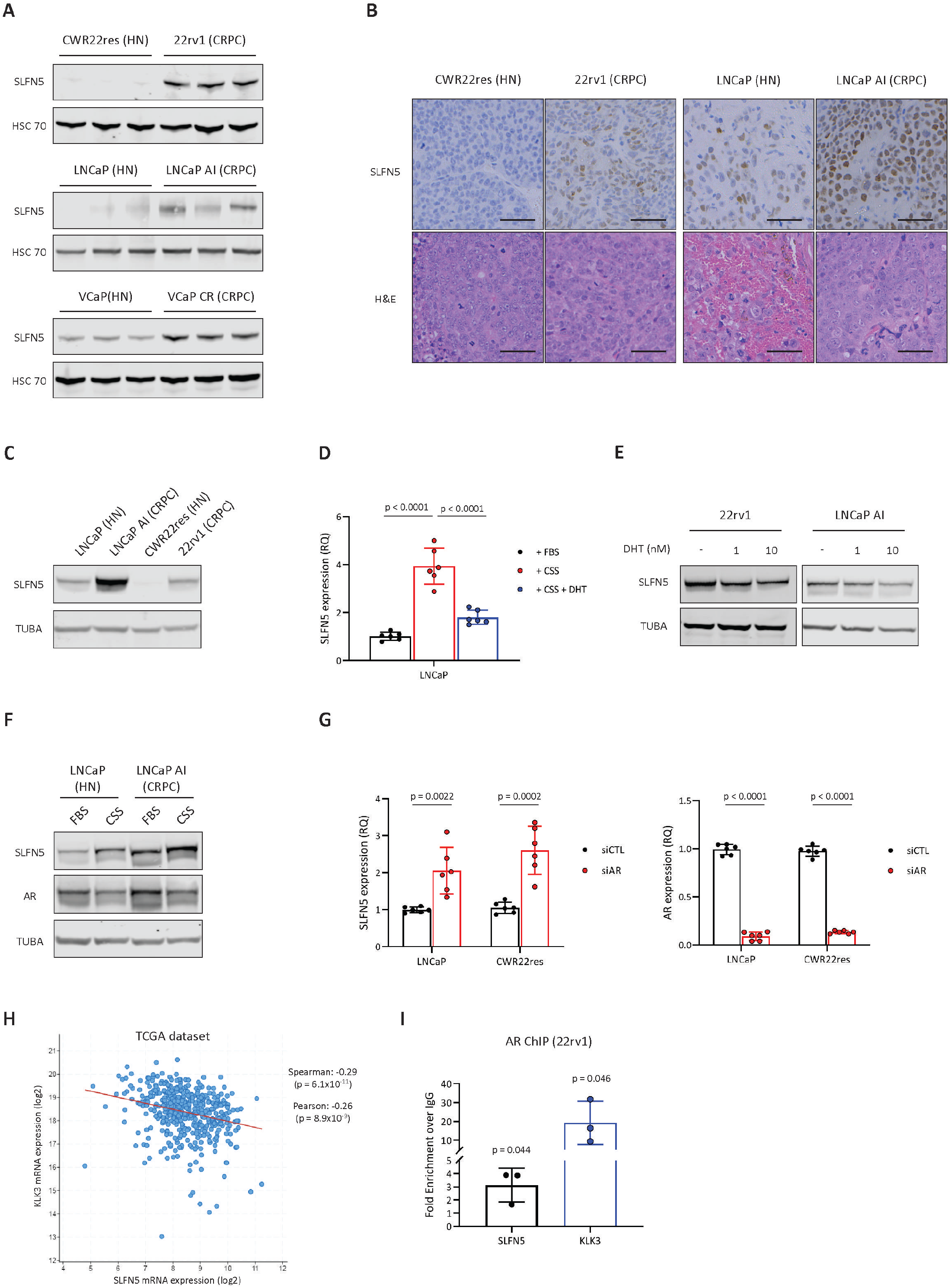
*SLFN5* is an AR-regulated gene. **A**, Western blot analysis of SLFN5 expression in HN and matched CRPC tumour orthografts. **B**, Immunohistochemical staining (top) of SLFN5 expression in HN (CWR22res and LNCaP) and matched CRPC (22rv1 and LNCaP AI) orthografts; and representative pictures of hematoxylin/eosin staining (bottom) of the corresponding orthografts. Scale bar represents 50 μm. **C**, Western blot analysis of SLFN5 expression in HN and matched CRPC cell lines. **D**, RT-qPCR analysis of *SLFN5* expression in LNCaP cells treated with DHT for 48 hours in androgen-depleted (CSS) conditions. **E**, Western blot analysis of SLFN5 expression in CRPC cell lines treated with DHT for 72 hours. **F**, Western blot analysis of SLFN5 and AR expression in LNCaP and LNCaP AI cells cultured in presence (FBS medium) or absence (CSS medium) of androgens for 72 hours. **G**, RT-qPCR analysis of *SLFN5* and *AR* expression in LNCaP and CWR22res cells silenced for AR expression. **H**, Correlation of *SLFN5* and *KLK3* mRNA levels in the PRAD TCGA dataset (cbioportal). **I**, RT-qPCR analysis of the *SLFN5* and *KLK3* promoters after anti-AR chromatin immunoprecipitation performed in 22rv1 cells. Panels **A, C, E, F**: HSC70 is used as a sample loading control. Panels **D, G**: *CASC3* was used as a normalising control. Panels **D, G, I**: Data are presented as mean values +/− SD. Panel **D**: *p-value using a 1-way ANOVA with a Tukey’s multiple comparisons test. Panel **G, I**: *p-value using a two-sided Student’s t-test.

### SLFN5 is associated with poor outcome in PCa patients

To assess the clinical relevance of SLFN5, we applied immunohistochemistry to assay for SLFN5 protein expression in a cohort of radical prostatectomy specimens. Similar to data from orthografts, SLFN5 immunoreactivity was primarily observed in the nuclei of epithelial cells (Figure 3a). SLFN5 expression was found to be highest in CRPC tumours (n = 45, p < 0.0001), followed by CRPC tumours with a neuroendocrine phenotype (NEPC, n = 29, p = 0.0165) in comparison to treatment naïve tumours (n = 151) (Figure 3b). Interestingly, compared to untreated tumour, SLFN5 expression was not altered following neo-adjuvant hormonal therapy (NHT-treated, median treatment time of 6 months, n = 162). High SLFN5 expression in patients significantly correlated with shorter relapse free survival (evaluated as time to biochemical relapse, p = 0.004, Figure 3c). Furthermore, SLFN5 expression was significantly elevated in high Gleason score tumours (>7 versus ≤7, p = 0.013) (Figure 3d) and was significantly associated with increased risk of metastasis (p-value = 0.0003) (Figure 3e).

**Figure 3:**
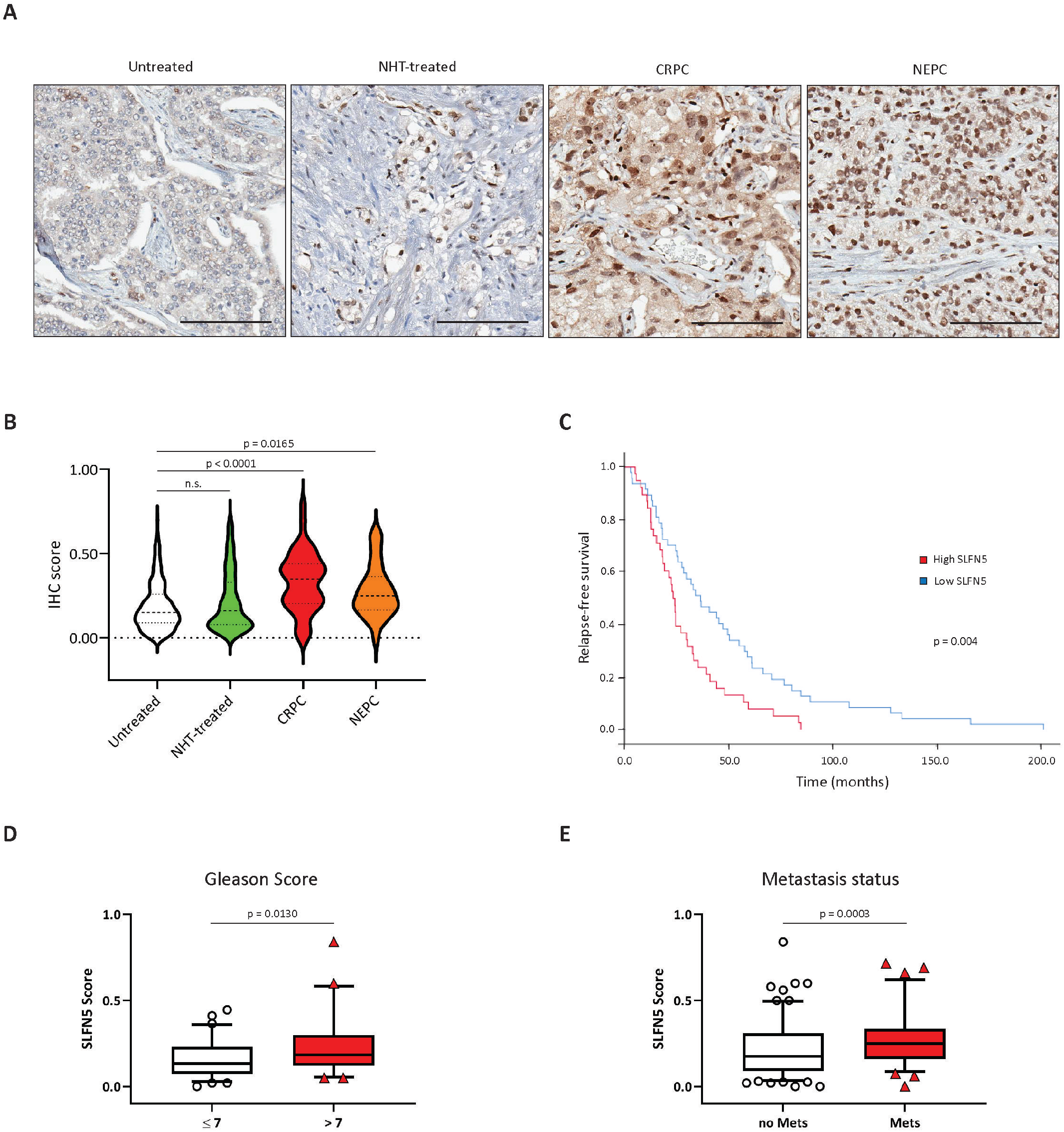
SLFN5 expression is high in CRPC tumours and correlates with poor patient outcome. **A**, Immunohistochemical staining of SLFN5 expression in treatment naïve, NHT-treated, CRPC and NEPC tumours. Scale bar represents 100 μm. **B**, Quantification of SLFN5 expression in PCa tissue samples. **C**, Kaplan-Meier relapse-free survival analysis of prostate cancer patients stratified according to median SLFN5 expression. Time to PSA recurrence was used as biochemical parameter. **D**, **E**, Quantification of SLFN5 expression in PCa tissue samples according to Gleason score and metastatic status. Centre line corresponds to median of data, top and bottom of box correspond to 95th and 5th percentile, respectively. Whiskers extend to adjacent values (minimum and maximum data points not considered outliers). Panel **B**: statistical analysis was performed using a 1-way ANOVA with a Dunnett’s multiple comparisons test. Panels **D**, **E**: statistical analysis was performed using a two-tailed Mann-Whitney U test. Panel **C**: statistical analysis was performed using a log rank test.

Taken together, these results suggest that SLFN5 has the potential to be used as a clinically relevant biomarker for aggressive PCa.

### SLFN5 loss impairs *in vivo* growth of CRPC tumours

To evaluate the functional importance of SLFN5 in CRPC, we used CRISPR-CAS9 technology to generate SLFN5 knockout (SLFN5 KO) clones in 22rv1 and LNCaP AI cells and assessed the proliferative and migratory abilities of CRPC cells *in vitro* (Figure 4a). While loss of SLFN5 did not consistently affect proliferation of 22rv1 cells (Figure 4b), both proliferation (Figure 4b) and migration (Figure 4c) were reduced in the LNCaP AI SLFN5 KO cells.

**Figure 4:**
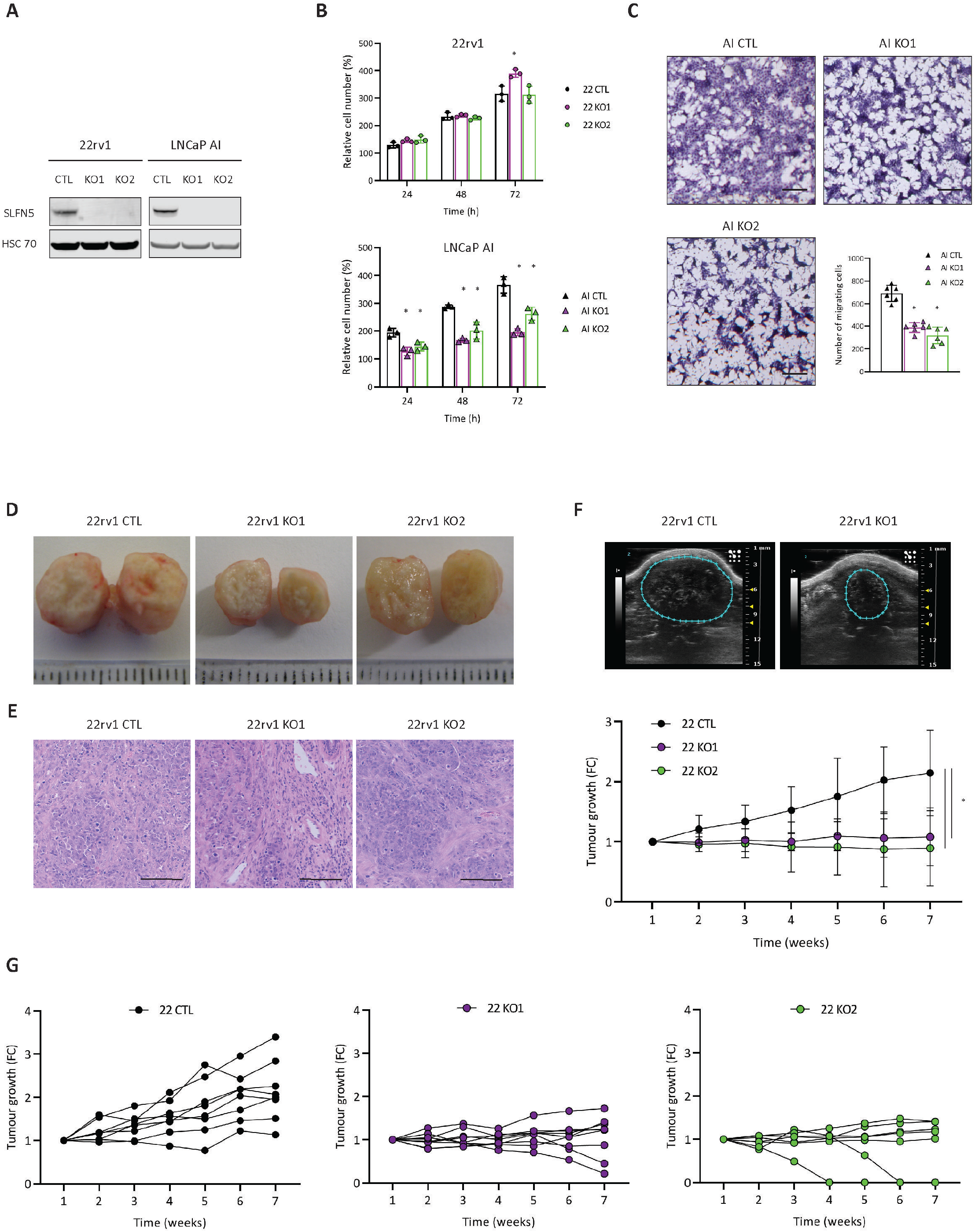
SLFN5 KO affects CRPC *in vitro* cell migration and *in vivo* tumour growth. **A**, Western blot analysis of SLFN5 expression in SLFN5 KO cells. HSC70 is used as a sample loading control. **B**, Cell proliferation of SLFN5 KO (knockout) and untargeted control (CTL) cells after 24, 48 and 72 hours. Cell count is normalised to initial number of cells at the start of the experiment. **C**, Cell migration of LNCaP AI SLFN5 KO (knockout) and untargeted control (CTL) cells after 48 hours. **D**, Representative pictures of 22rv1-derived SLFN5 KO (knockout) and untargeted control (CTL) tumour orthografts. **E**, Representative pictures of hematoxylin/eosin staining of the corresponding tumour orthografts. Scale bar represents 100 μm. **F**, Representative pictures of 22rv1 CTL or SLFN5 KO tumour orthografts monitored by ultrasound imaging (top). Quantification of tumour volume along time using ultrasonography (bottom). **G**, same as **F** but individual tumours are plotted on separate graphs. Panels **B, C, F**: Data are presented as mean values +/− SD. Panels **B, C**: *p-value < 0.05 using a 1-way ANOVA with a Dunnett’s multiple comparisons test. Panel **F**: *p-value < 0.05 using a 2-way ANOVA with a Dunnett’s multiple comparisons test.

Because SLFN5 was originally discovered in an *in vivo* proteomic screen, we speculated whether SLFN5 could affect the growth of CRPC tumours *in vivo*. To test this hypothesis, 22rv1 control and SLFN5 KO (CTL and KO respectively) cells were orthotopically injected into castrated mice and tumour volume was monitored weekly using ultrasonography. While SLFN5-deficient cells remained able to form solid tumours in CRPC condition (Figure 4d,e), tumour growth was strongly reduced in absence of SLFN5 (Figure 4f). In addition, partial or total tumour regression was also observed in around 25% of SLFN5 KO tumours (Figure 4g). Taken together, these results suggest that SLFN5 is important for tumour adaptation to CRPC condition, rather than for tumour initiation, and inhibiting SLFN5 may present as a potential target to regress some tumours.

### SLFN5 depletion remodels the transcriptome of CRPC cells

SLFN5 has been described as a transcriptional modulator in glioblastoma^19^. In agreement with a potential role in regulating transcriptional activity, SLFN5 was expressed in the nucleus of CRPC cells (Figure 5a). To understand the molecular functions of SLFN5, we compared the transcriptome of SLFN5 KO and CTL 22rv1 cells (Supplementary Data 2). Loss of SLFN5 resulted in significant alteration of 428 genes (FC = 2; p-value < 0.05), with 331 up-regulated and 97 down-regulated genes in the same direction in both KO clones when compared to CTL cells (Figure 5b). Enrichment Pathway Analysis emphasised that many transcripts altered in SLFN5 KO cells encoded for plasma membrane proteins. Cell adhesion was one of the top up-regulated pathways (FC > 2), while cell locomotion/migration, extracellular matrix organisation and ion transport were among the pathways that were significantly reduced in KO cells (FC < −2) (Figure 5c and Supplementary Data 3). We performed a similar RNAseq analysis on SLFN5-proficient and -depleted 22rv1 orthografts. Even with higher variability among *in vivo* tumour samples, we observed 88 genes that were significantly modulated in SLFN5-deficient tumours (FC = 2, p-value < 0.05), with the majority (n=68) of these genes being down-regulated in KO tumours (Figure 5b, Supplementary Data 2). Importantly, 22 genes were strongly down-regulated (FC < −3) in absence of SLFN5 in both the *in vitro* and *in vivo* analyses. This allowed us to define a signature of SLFN5-target genes in CRPC (Table 1). Finally, RNAseq expression data for the top down-regulated candidates were validated by qPCR (Figure 5d).

**Figure 5:**
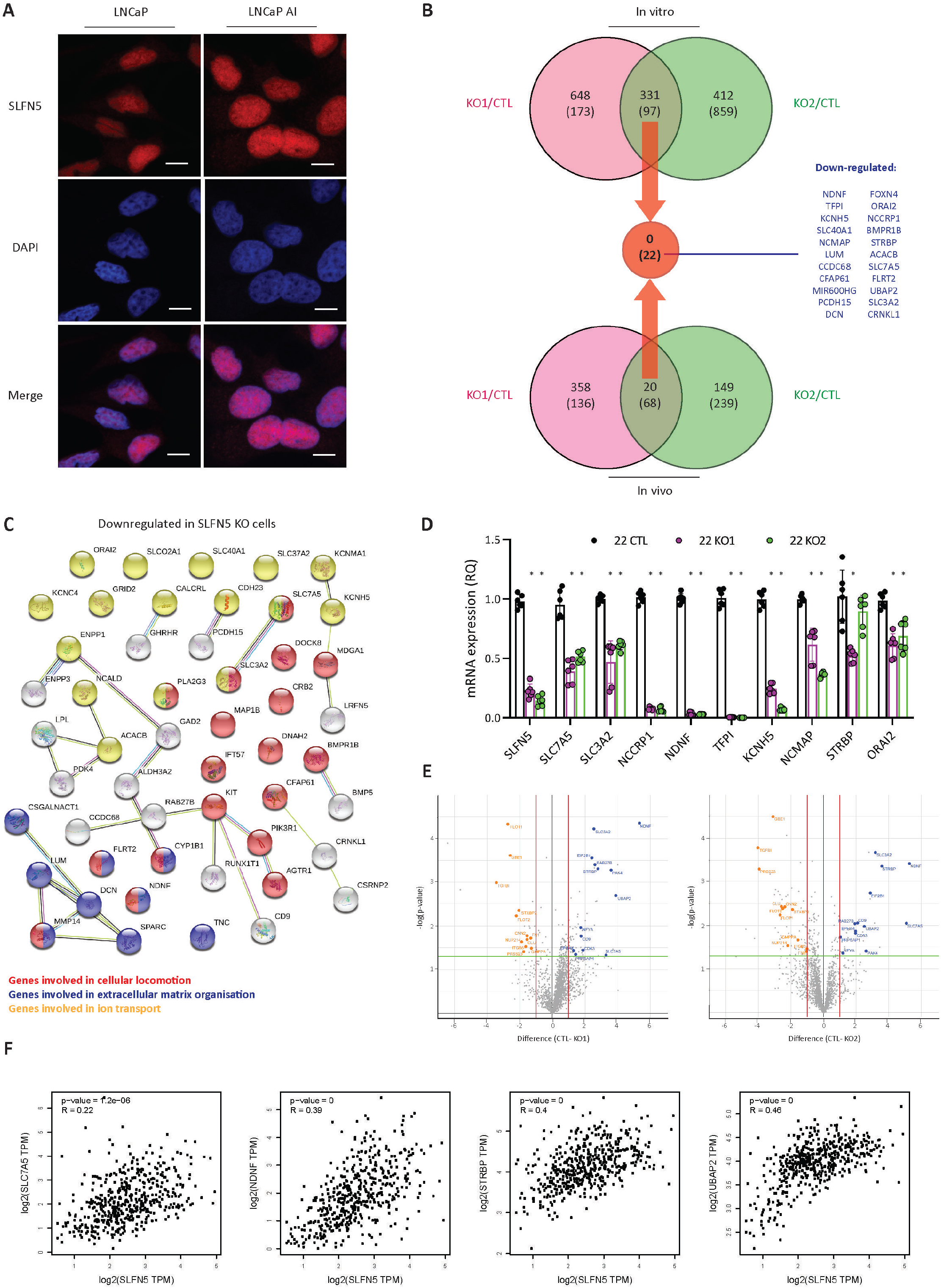
SLFN5 KO remodels the transcriptome of CRPC cells. **A**, Immunofluorescence showing nuclear SLFN5 expression in LNCaP and LNCaP AI cells. Scale bar represents 10 μm. **B**, Venn diagrams highlighting genes commonly modulated (p-value < 0.05, FC = 2) in SLFN5 KO cells (top) and tumours (bottom) when compared to their respective controls. Up-regulated genes are on top; Down-regulated genes are into brackets. **C**, Schematic representation of the down-regulated genes (p-value < 0.05, FC = 2) in SLFN5 KO cells when compared to CTL cells. Pathway enrichment analysis was performed using the STRING database (http://string-db.org). **D**, RT-qPCR analysis of *SLFN5* and top downregulated genes from **B** in SLFN5 KO and CTL cells. **E**, Volcano plots showing the proteins significantly modulated (p-value < 0.05, FC = 2) in the proteomic analysis of SLFN5 KO tumours. **F**, Pearson’s correlation analysis of *NDNF*, *STRBP, UBAP2* and *SLC7A5* with *SLFN5* using the PRAD TCGA dataset. Results were obtained using the GEPIA website http://gepia.cancer-pku.cn/. Panel **D**: Data are presented as mean values +/− SD. Panel **D**: *p-value < 0.05 using a 1-way ANOVA with a Dunnett’s multiple comparisons test. Panel **F**: statistical analysis was performed using a logrank test.

To assess whether the transcriptional changes occurring in the absence of SLFN5 would reflect at the protein level, we further performed a proteomic comparison of the SLFN5 KO and CTL orthografts. The analysis highlighted 25 proteins that were significantly modulated (FC = 2, p-value < 0.05) in the absence of SLFN5 (Figure 5e). Among these candidates, 5 proteins (NDNF, STRBP, UBAP2, SLC7A5 and SLC3A2) belonged to the SLFN5-gene signature defined in Table 1. Moreover *NDNF*, *STRBP*, *UBAP2* and *SLC7A5* transcript levels strongly correlated with SLFN5 expression in PCa patients (Figure 5f, TCGA dataset), thus supporting a potential transcriptional regulation by SLFN5.

Of note, SLC7A5 and SLC3A2 are the functional components of the AA transporter LAT1 which has recently gained interest as a molecular target for cancer therapies^22^. We therefore sought to explore the link between SLFN5 and LAT1 in PCa.

### SLFN5 controls LAT1 expression in CRPC

Down-regulation of the LAT1 (SLC7A5/SLC3A2) complex was confirmed in 22rv1-derived SLFN5 KO cells and orthografts (Figure 6a-b). Importantly, SLC7A5 expression was also decreased in LNCaP AI cells depleted for SLFN5 (Figure 6c), and increased in SLFN5-overexpressing LNCaP (Figure 6d). Overall, SLC7A5 protein level was high in CRPC cell lines when compared to HN cells, and positively correlated with SLFN5 expression (Figure 6e).

**Figure 6:**
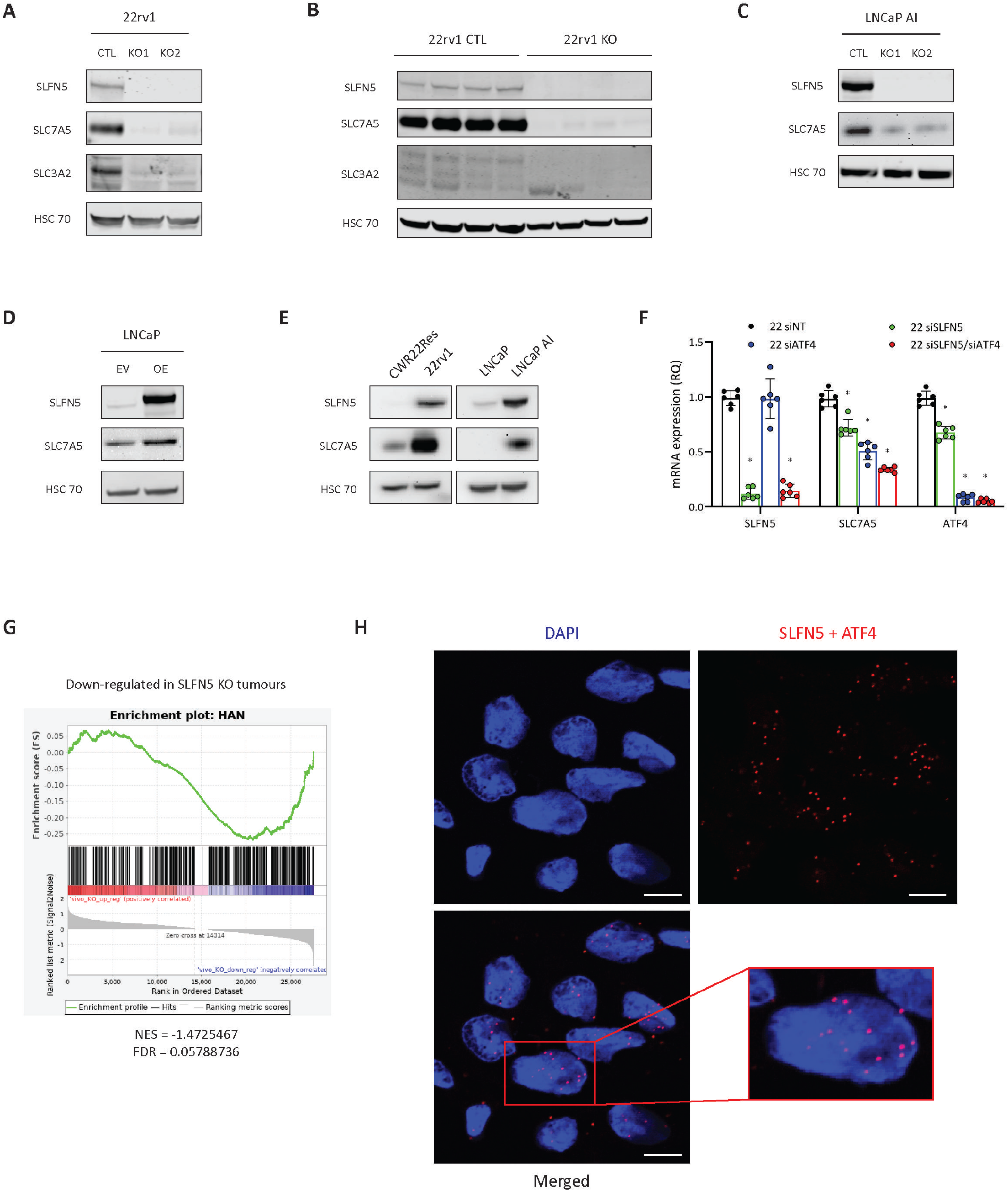
SLFN5 regulates LAT1 expression in CRPC. **A**, Western blot analysis of SLFN5, SLC7A5 and SLC3A2 expression in 22rv1 SLFN5 KO and CTL cells. **B**, Western blot analysis of SLFN5, SLC7A5 and SLC3A2 expression in 22rv1-derived SLFN5 KO and CTL tumour orthografts. **C**, Western blot analysis of SLFN5 and SLC7A5 expression in LNCaP AI SLFN5 KO and CTL cells. **D**, Western blot analysis of SLFN5 and SLC7A5 expression in LNCaP cells overexpressing SLFN5. **E**, Western blot analysis of SLFN5 and SLC7A5 expression in HN and matched CRPC cell lines. **F**, RT-qPCR analysis of *SLFN5*, *SLC7A5* and *ATF4* expression in 22rv1 cells silenced for SLFN5, ATF4 or both. **G**, Gene set enrichment plots analysed from SLFN5-depleted tumours transcriptomics using ATF4-related gene set obtained from^59^. **H**, Proximity ligation assay of SLFN5 and ATF4 performed on 22rv1 cells. Red dots represent co-localisation. Scale bar represents 11 μm. Panels **A**, **B**, **C**, **D**, **E**: HSC70 is used as a sample loading control. Panel **F**: Data are presented as mean values +/− SD. Panel **F**: *p-value < 0.05 using a 1-way ANOVA with a Tukey’s multiple comparisons test.

We next investigated the mechanism by which SLFN5 regulates LAT1 in CRPC. In a recent study, ChIPseq analysis following SLFN5 immunoprecipitation revealed the presence of SLFN5-specific binding motifs in the genome of breast cancer cells^18^. Interestingly, both *SLC7A5* and *SLC3A2*, as well as *NDNF* and *STRBP*, displayed enrichment of SLFN5-binding motifs in their promoters (Supplementary data 4). However, transient silencing of SLFN5 by siRNA only moderately reduced *SLC7A5* expression (Figure 6f). *SLC7A5* and *SLC3A2* have also been reported as ATF4 target genes in cancer^13^. In addition, negative enrichment of several ATF4-related genesets was observed in the transcriptomic analysis of the SLFN5 KO tumours (Figure 6g and Supplementary figure 2a), and majority of the 22 SLFN5-regulated genes (Figure 5b, Table 1) were predicted to harbour strong binding sites for ATF4 (Supplementary data 5). We therefore hypothesized that SLFN5 could act as a transcriptional modulator of ATF4 in PCa. Proximity ligation assay performed on 22rv1 cells confirmed a direct interaction between SLFN5 and ATF4 (Figure 6h). Moreover, enrichment of the SLFN5-specific motif was observed in the *ATF4* promoter (Supplementary data 4) and SLFN5 depletion indeed reduced *ATF4* mRNA expression (Figure 6f). Silencing of ATF4 in 22rv1 cells also led to a strong decrease in *SLC7A5* mRNA expression, and this effect was amplified when SLFN5 and ATF4 were both silenced concomitantly (Figure 6f). Finally, co-silencing of SLFN5 and ATF4 also reduced the expression of other genes within the SLFN5 signature, such as *KCNH5*, *NCMAP* and *NCCRP1* (Supplementary Figure 2b). Taken together, these results suggest a role for SLFN5 in the ATF4-mediated regulation of LAT1 in CRPC.

### SLFN5 drives LAT1-mediated activation of mTOR in CRPC

LAT1 is a large neutral AA transporter that controls the cellular uptake of branched chain and aromatic AAs in exchange of glutamine^10^. Hence, LAT1 expression is crucial for the regulation of cancer cell metabolism. To evaluate the impact of SLFN5-LAT1 depletion on the metabolism of CRPC cells, we compared the metabolic profiles of 22rv1 CTL and SLFN5 KO cells using LC-MS metabolomics. Consistent with the role of LAT1 in AA homeostasis, we observed that the levels of many AAs (Lys, Arg, Orn, Met, Leu, Ile, Tyr) were decreased in the SLFN5 KO cells (Figure 7a). By contrast, SLFN5-deficient cells also showed elevated levels of glutathione, in both reduced and oxidised forms (Figure 7a).

**Figure 7:**
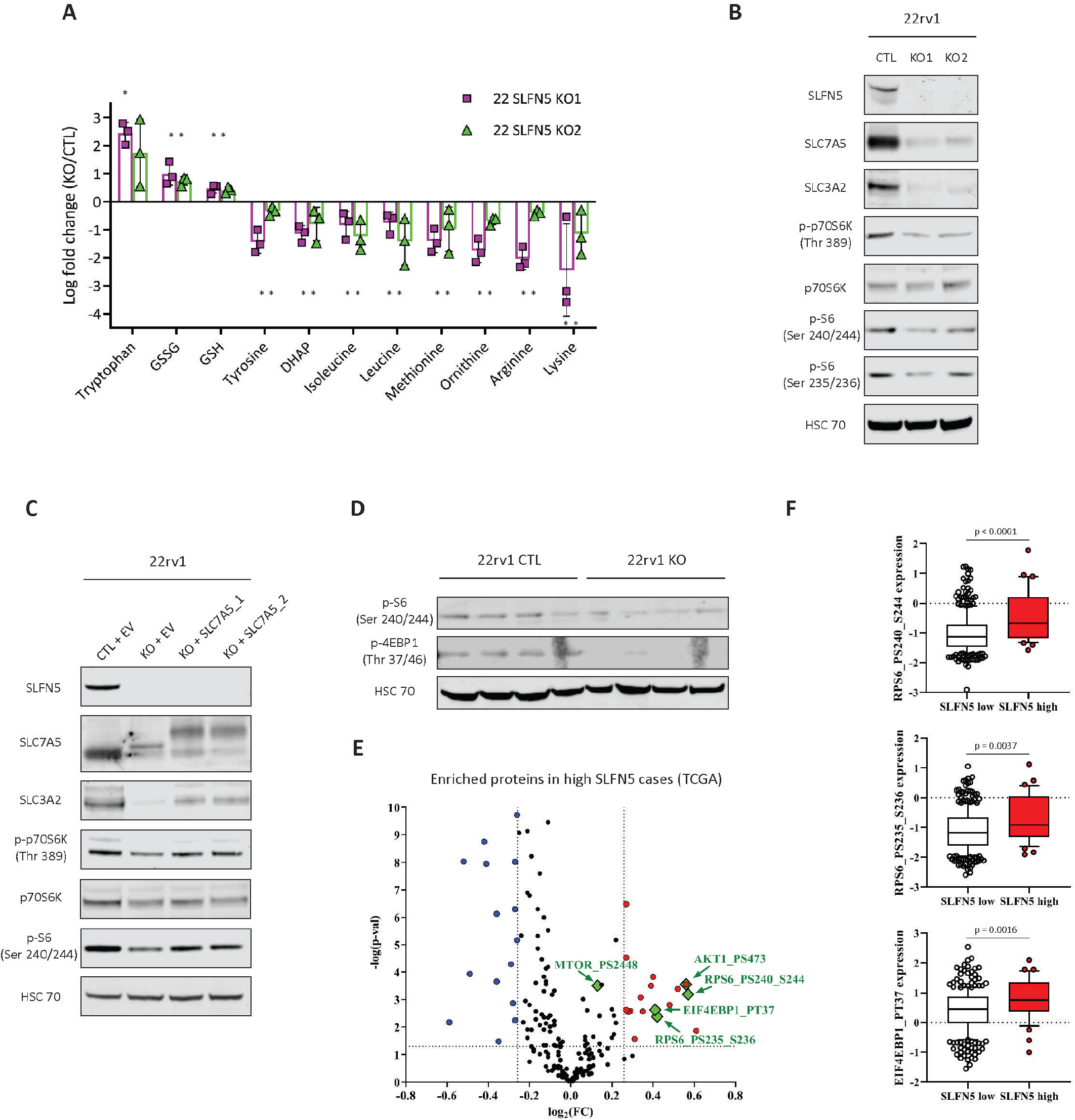
SLFN5 expression promotes LAT1-dependent activation of mTORC1 signalling in CRPC. **A**, Steady-state levels of significantly regulated metabolites in SLFN5 KO cells when compared to CTL cells (FC > 1.2). Selected metabolites were significantly altered in at least one of the two KO cells (p < 0.05 using two-sided Student’s t-test). **B**, Western blot analysis of SLFN5, SLC7A5, SLC3A2, p-70S6K and p-S6 expression in 22rv1 SLFN5 KO and CTL cells. **C**, same analysis as **B** performed on 22rv1 SLFN5 KO and CTL cells overexpressing two different SLC7A5 constructs. **D**, Western blot analysis of p-S6 and p-4EBP1 expression in 22rv1-derived SLFN5 KO and CTL tumour orthografts. **E**, Volcano plot of the modulated proteins between high and low SLFN5 tumours using the TCGA PRAD dataset. Red and blue dots represent the proteins that are significantly up-regulated and down-regulated (FC = 1.2, p < 0.05) in high SLFN5 tumours respectively. **F**, Differential expression of p-S6 (Ser235-236 and 240-244) and p-4EBP1 (Thr37) in high and low SLFN5 tumours, according to the data generated in **E**. Centre line corresponds to median of data, top and bottom of box correspond to 90th and 10th percentile, respectively. Whiskers extend to adjacent values (minimum and maximum data points not considered outliers). Panels **B**, **C**, **D**: HSC70 is used as a sample loading control. Panel **A**: Data are presented as mean values +/− SD. Panel **F**: statistical analysis was performed using a two-tailed Mann-Whitney U test.

Changes in amino acid homeostasis are known to regulate the mTOR signalling pathway^23^. Consistent with the observed changes in AA levels, SLFN5-deficient cells exhibited impaired mTOR activity, as evidenced by decreased phosphorylation levels of S6K and S6 proteins, which are two downstream targets of mTORC1 (Figure 7b). To test whether mTOR activation was dependent on LAT1, we stably re-expressed a myc-tagged version of SLC7A5 in SLFN5-deficient cells (Figure 7c). Reintroducing SLC7A5 in SLFN5 KO cells was sufficient to restore the phosphorylation levels of S6K and S6, therefore indicating that impaired mTOR signalling in SLFN5-deficient cells was due, at least in part, to the decrease in LAT1 expression. Importantly, impaired mTOR signalling was also observed *in vivo* in SLFN5 KO orthografts (Figure 7d). Finally, using the TCGA PRAD dataset we identified proteins that were significantly modulated in high vs. low SLFN5 tumours (Figure 7e). Strikingly, several down-stream effectors of the mTOR signalling pathway (p-AKT Ser473, p-S6 Ser235-236 and Ser240-244, p-4EBP1 Thr37 and p-mTOR Ser2248) were positively enriched in high SLFN5 tumours. Taken together, these results suggest SLFN5 as a novel regulator of mTOR signalling in PCa.

## Discussion

Overcoming resistance to AR targeted therapies remains the ultimate goal for the treatment of advanced PCa. CRPC develops in the majority of patients treated with ADT and is often associated with metastasis. The molecular heterogeneity of the CRPC disease reflects the numerous ways that tumours can evolve to escape current therapies. Indeed, point mutations^24,25^, genomic deletion^26^ or amplification^27^ of the AR gene, reprogramming of AR signalling^28^ as well as compensations from other signalling pathways^26,29^ can all account for resistance to ADT^30^. In this study, we characterised three different *in vivo* models that were generated to specifically study CRPC and ADT resistance. These models consist of the orthotopic injection of three pairs of isogenic, AR-responsive, human cancer cell lines into the prostate of immuno-deficient mice, before undergoing orchidectomy to achieve ADT. In depth proteomic characterisation of these three models highlighted pathways and molecular markers that are commonly involved in CRPC. Despite molecular differences between the models, that is reminiscent of the molecular heterogeneity observed in clinical CRPC samples^30^, our analysis indicated that resistance to ADT was accompanied with a major rewiring of tumour metabolism, especially lipid and AA metabolism. In addition to steroid biogenesis, which plays an important role in CRPC development^31^, branched chain amino acid (BCAA) and fatty acid (FA) degradation were strongly dysregulated upon resistance to ADT. BCAA catabolism serves to replenish the tricarboxylic acid cycle and is dysregulated in PCa^32^, while targeting FA metabolism has been proposed as a therapeutic option in the context of CRPC and enzalutamide resistance^33,34^. Ferroptosis, another lipid-related process that has recently been suggested as an important resistance mechanism against AR targeted therapies^6^, was also enriched in our CRPC models. Likewise, peroxisome proliferator-activated receptor (PPAR) signalling was commonly enriched in all three models of CRPC orthografts. While AR itself remains the most influential regulator of PCa metabolism^5^, the contribution of PPAR signalling pathways to the regulation of PCa lipid metabolism has recently gained interest. For example, PPARγ has been identified as a critical regulator of PCa invasion and metastasis^35^, while PPARα is an established AR target gene that is overexpressed in advanced PCa^36^.

As well as uncovering the molecular pathways that are associated with CRPC, our analysis allowed us to define a repertoire of several candidates whose expressions were robustly associated with CRPC. Among these proteins, we focused our attention on Schlafen Family Member 5, a potential transcription co-regulator whose expression had not yet been reported in PCa. SLFN5 expression was increased in CRPC patients and was associated with poor clinical outcome. This increase in expression can be explained by the AR-dependent regulation of SLFN5 in prostate cells, an observation that was supported by the presence of AR-binding sites in the promoter region of SLFN5^37^. In line with a pro-malignant role of SLFN5 in PCa, CRISPR-mediated KO of SLFN5 in CRPC cells reduced cell migration and further impaired CRPC tumour growth in castrated mice. The role of SLFN5 in cancer remains controversial. In contrast to our data, early studies in melanoma^15^, breast^16^ and renal cell carcinoma^17^ suggested that high SLFN5 expression was correlated with favourable patient outcomes. Moreover, in these cell types, SLFN5 silencing resulted in enhanced cell migration and invasion, thus suggesting a tumour suppressive role for SLFN5. Conversely, pro-tumoural properties of SLFN5 have been described in glioblastoma (GBM), along with a transcriptomic-informed signature of SLFN5 genes in U87 glioblastoma cells^19^. Surprisingly, there was little overlap between the SLFN5 gene signature that we identified in CRPC cells and that reported for U87 cells, raising the possibility that SLFN5-mediated transcriptional activities might be cell type and/or context dependent.

By combining transcriptomics and proteomics, we identified SLC7A5 and SLC3A2, the two components of the LAT1 amino acid transporter, as targets of SLFN5 in CRPC. LAT1 is a member of the system L transporter family and mediates the intracellular uptake of BCAA and aromatic AA in exchange for glutamine^11^. In PCa, LAT1 expression is associated with an increased risk of metastasis^13^. LAT1 is also up-regulated following androgen deprivation^12,13^ and is an independent predictor of castration resistance^12^. Moreover targeting LAT-dependent AA transport has shown promising results in preclinical models^22^. In CRPC cells, SLFN5 KO led to a strong downregulation of the LAT1 transporter, although the molecular mechanisms underlying this observation remain to be fully uncovered.

Based on our observation that transient SLFN5 silencing only moderately reduced SLC7A5 expression, we hypothesised that SLFN5 co-regulated SLC7A5 expression along with another transcription factor. An interesting candidate is the stress-induced factor ATF4, which has been implicated in CRPC^38^. Indeed, SLC7A5 is an established target of ATF4 in PCa^13^, and we have shown that SLFN5 and ATF4 were able to physically interact with each other. Moreover, the majority of the SLFN5-regulated genes (Table 1) displayed strong ATF4-binding sites, and both ATF4 and SLC7A5 also presented SLFN5-enriched motifs in their promoter region. Finally, co-silencing of SLFN5 and ATF4 dramatically reduced the expression of the LAT1 transporter. Additional research is required to further define the molecular mechanisms connecting SLFN5 and ATF4 in the context of CRPC.

Consistent with a role for LAT1 in maintaining AA homeostasis, we observed that SLFN5 KO cells displayed low intracellular levels of essential AA, which are potent activators of mTORC1 signalling. Leucine for example is important to maintain mTORC1 localisation at the lysosomal surface, subsequently allowing activation of the downstream signalling pathway^9^. Therefore a role for LAT1 in stimulating mTORC1 signalling has been reported in the pathology of multiple diseases^11^. mTOR is also frequently dysregulated in advanced prostate cancer^7^ and targeting this pathway has shown promising results in preclinical models^39^. However, mTOR inhibitors have shown limited efficacy in the clinic^8^. One reason could be that only a subpopulation of PCa patients might benefit from such mTOR-targeted therapies. Interestingly, using publicly available data, we showed that high SLFN5 levels in patients correlate with increased phosphorylation levels of multiple down-stream effectors of the mTOR signalling pathway (p-mTOR, p-S6 and p-4EBP1). Therefore, SLFN5 expression, as well as the identification of a CRPC specific SLFN5-gene signature, could help in stratifying patients that would benefit from mTOR inhibition and may present as a potential therapeutic candidate to resensitize patients to treatment.

In conclusion, this study provides an in-depth characterisation of innovative preclinical models of CRPC that were generated to recapitulate specific molecular features of ADT resistance in PCa. Our results confirm the suitability of these orthograft models to account for the high degree of heterogeneity observed in CRPC patients, and further highlight the transcriptional modulator SLFN5 as a clinically relevant target for CRPC. Mechanistically, SLFN5 controls the expression of the LAT1 transporter in CRPC cells, potentially acting through an ATF4-dependent mechanism. As a result SLFN5 deletion impairs CRPC tumour growth *in vivo*, alters CRPC cell metabolism and disrupts mTORC1 signalling in a LAT1-dependent manner. Taken together, our results support the idea of targeting metabolism for the treatment of PCa, and further establish SLFN5 as a potential target and an important metabolic regulator in CRPC.

## Methods

### Cell culture

Hormone naïve cells (CWR22res, LNCaP and VCaP) were cultured in RPMI medium (Gibco, Thermo Fisher Scientific, Waltham, MA, USA) supplemented with 10% foetal bovine serum (FBS, Gibco, Thermo Fisher Scientific, Waltham, MA, USA) and 2 mM glutamine (Gibco, Thermo Fisher Scientific, Waltham, MA, USA). Castration-resistant cells (22rv1 and LNCaP AI) were cultured in androgen-deprived medium consisting of phenol-free RPMI (Gibco, Thermo Fisher Scientific, Waltham, MA, USA) supplemented with 10% charcoal stripped serum (CSS, Gibco, Thermo Fisher Scientific, Waltham, MA, USA) and 2 mM glutamine. All cells were kept in incubators set at 37°C and 5% CO2. LNCaP (ATCC CRL-1740), 22Rv1 (ATCC CRL-2505), and VCAP (ATCC CRL-2876) were obtained from ATCC. CWR22Res cells (hormone-responsive variant of CWR22 cells) were obtained from Case Western Reserve University, Cleveland, Ohio. LNCaP AI cells were obtained from Newcastle University, UK.

### Generation of stable knockout and overexpressing cells

All plasmids were transfected into 10^6^ cells using Cell Line Nucleofector Kit V (Lonza, Basel, Switzerland) and the T-013 program of a Nucleofector 2b Device (Lonza, Basel, Switzerland).

22rv1 and LNCaP AI cells were transfected with commercially available *SLFN5* CRISPR/Cas9 KO Plasmid (sc-408333) and *SLFN5* HDR Plasmid (sc-408333-HDR) or control plasmids (Santa Cruz Technologies, Dallas, TX, USA). Cells were then put under clonal selection to generate single-cell colonies. CTL and KO clones were then expanded and further selected for experiments based on SLFN5 protein expression.

For overexpressing cells, LNCaP cells were transfected with *SLFN5* (NM_144975) Human MYC-Tagged ORF Clone (RC216330, Origene, Rockville, MD, USA) or the corresponding empty vector plasmid (PS100001, Origene, Rockville, MD, USA). 22Rv1 SLFN5 KO cells were further transfected with *SLC7A5* (NM_003486) Human Tagged ORF Clone (RC207604, Origene, Rockville, MD, USA). Cells were then clonally selected, expanded and ultimately selected for further experiments based on SLFN5 or SLC7A5 protein expression.

### siRNA transfection

750.000 cells were seeded in 6-well plates and allowed to attach overnight. The next day, transfections were performed using Lipofectamine RNAimax (Invitrogen, Thermo Fisher Scientific, Waltham, MA, USA) according to the manufacturer’s protocol. ON-TARGETplus smartpool siRNAs against AR (L-003400-00) and ATF4 (L-005125-00), as well as non-targeting siRNA (D-001810-01-20) were purchased from Dharmacon (Dharmacon, Horizon inspired cell solutions, Cambridge, UK). RNA or protein extraction was performed 72 hours after transfection.

### Chromatin immunoprecipitation (ChIP)

Chromatin was prepared with the truChIP™ Chromatin Shearing Kit (Covaris, Brighton, UK) according to manufacturer’s instructions. Each sample was sonicated for 10 min using Covaris sonicator. ChIP were performed using the IP-Star Compact Automated System (Diagenode, Liege, Belgium). Four μg of isolated chromatin was immunoprecipitated with either 1 μg ChIP grade antibody (anti-AR 17-10489, Millipore Burlington, MA, USA) or 1μg of IgG (C15410206, Diagenode, Liege, Belgium) in dilution buffer (0.01% SDS; 1.1% Triton X 100; 2 mM EDTA; 16.7 mM Tris-Cl pH 8.0; 167 mM NaCl; 1× protease inhibitor cocktail, Sigma Aldrich, St Louis, MI, USA). The DNA/protein complexes were washed four times in IP Wash buffer (100 mM Tris-HCl pH 8.0; 500 mM LiCl 1%; Triton X100; 1% Deoxycholic Acid. After reversal of crosslinking, the immunoprecipitated DNA was purified by a regular DNA extraction protocol and analysed employing RT-qPCR with the SYBR-Green Takara (Ozyme, Paris, France) and step one plus applied Real-Time PCR system. The PCR conditions were 10 min at 95 °C followed by 45 cycles of 10 s at 95 °C, 30 s at 60 °C, and 30 s at 72 °C.

### Cell proliferation

CTL or SLFN5 KO cells were seeded in 24-well plates (70.000 cells per well) and allowed to attach for 24 hours. Cells were then immediately counted using a CellDrop Cell Counter (DeNovix, Wilmington, DE, USA) (= T0) or allowed to grow for an additional 72 hours following medium replacement. Cells were then counted at 24, 48 and 72 hours after medium change. Cell number was normalised to the initial cell count at T0.

### Cell migration

LNCaP AI cells were kept in serum-free medium for 24 hours before the experiment. Next, 500 μl of FBS-supplemented medium were dispensed in a 24-well plate. Cells were then trypsinised and resuspended in serum-free medium at a concentration of 10^6^ cells/ml, and 500 μl containing 500,000 cells were added on top of 8 μm pores inserts (Corning, New York, MA, USA). After 48 hours, the inserts were fixed in 100% methanol for 30 minutes at −20°C and subsequently stained with hematoxylin for 30 minutes at room temperature. Insert membranes were then washed with tap water and cells on the upper side were scrapped with a wet cotton bud. Finally, membranes were cut from the insert and mounted onto microscopy slides. Images were taken with a Zeiss AXIO microscope (Zeiss, Oberkochen, Germany) and further quantified using ImageJ software (v. 1.46r).

### Human CRPC orhografts

*In vivo* experiments were performed in accordance with the ARRIVE guidelines^40^, and were reviewed by a local ethics committee under the Project Licence P5EE22AEE in full compliance with the UK Home Office regulations (UK Animals (Scientific Procedures) Act 1986). Prostate cancer cells were suspended in serum-free RPMI medium and mixed 1:1 with Matrigel (Corning, NY, USA). Briefly, 14×10^6^ cells (in 50 μl) were injected into the anterior prostate of CD1-nude mice (Charles River Laboratories, Wilmington, MA, USA). For CRPC conditions, orchidectomy was performed at the time of injection. Tumour growth was monitored weekly using A Vevo3100 ultrasound imaging system (Fujifilm Visualsonics, The Netherlands). Tumours were then allowed to grow for 9 weeks before reaching endpoint. At the end of the experiment, tumour orthografts were collected and weighted. Half of the tumour material was fixed in 10% formalin for histological procedures and the other half was snap-frozen in liquid nitrogen for protein, mRNA, and metabolite extractions.

### Proteomic analysis of paired HN and CRPC orthografts

2-5 mg of tumour powder were resuspended in 150 μl of 4% SDS containing protease and phosphatase inhibitors. Samples were then sonicated and centrifuged at 16000 × g for 10 minutes. Supernatant was collected and quantified using the BCA protein assay kit (Thermo Fisher Scientific, Waltham, MA, USA). To prepare the super-SILAC standard, equal amounts of SILAC-labelled cell lysates from CWR22res, LNCAP, LNCAPAI, and VCAP cell lines were mixed together. The lysate obtained from each tumour sample was mixed at 1:1 ratio with the super-SILAC standard prior to FASP digestion^41^ with Endoproteinase Lys-C (Alpha Laboratories, UK) and trypsin (Promega, Madison, WI, USA).

Approximately 500 μg of SILAC-labelled protein digests were fractionated using high pH reverse phase chromatography. A C18 column (250 × 4.6 mm i.d. – Durashell RP (5 μm, 150Å)) was used with an HPLC system (Ultimate LPG-3000 binary pump and UVD170U Ultraviolet detector, Dionex, Thermo Fisher Scientific, Waltham, MA, USA). Modules were controlled by Chromeleon v. 6.7. Solvent A (98% water, 2% acetonitrile) and solvent B (90% acetonitrile and 10% water) were adjusted to pH 10 using ammonium hydroxide. Samples were injected manually through a Rheodyne valve onto the RP-HPLC column equilibrated with 4% solvent B and kept at this percentage for 6 minutes. A two-step gradient was applied at a flow-rate of 1 ml/min (from 4–28% B in 36 minutes, then from 28-50% B in 8 minutes) followed by a 5-minutes washing step at 80% solvent B and a 10-minutes re-equilibration step, for a total run time of 65 minutes. Column eluate was monitored at 220 and 280 nm and collected using a Foxy Jr. FC144 fraction collector (Dionex, Thermo Fisher Scientific, Waltham, MA, USA). Collection was allowed from 8 to 50 minutes for 85 seconds per vial for a total of 30 fractions. The first 4 and the last 5 fractions were pooled resulting in 21 fractions in total.

Each of the 21 fractions was dried down and then re-suspended in 2% acetonitrile/0.1% TFA acid in water and separated by nanoscale C18 reverse-phase liquid chromatography performed on an EASY-nLC II (Thermo Fisher Scientific, Waltham, MA, USA) coupled to a Linear Trap Quadrupole Orbitrap Velos mass spectrometer (Thermo Fisher Scientific, Waltham, MA, USA). Elution was carried out using a binary gradient with buffer A (2% acetonitrile) and B (80% acetonitrile), both containing 0.1% of formic acid. Peptides were subsequently eluted into a 20 cm fused silica emitter (New Objective, Woburn, MA, USA) packed in-house with ReproSil-Pur C18-AQ, 1.9 μm resin (Dr Maisch GmbH, Ammerbuch, Germany). The emitter was kept at 35°C by means of a column oven integrated into the nanoelectrospray ion source (Sonation, Germany). Peptides separation was performed using 3 different gradients optimised for different set of fractions as described previously^42^. Eluting peptides were electrosprayed into the mass spectrometer using a nanoelectrospray ion source (Thermo Fisher Scientific, Waltham, MA, USA). An Active Background Ion Reduction Device (ABIRD, ESI source solutions, Woburn, MA, USA) was used to decrease air contaminants signal level.

The mass spectrometer was operated in positive ion mode and used in data-dependent acquisition mode (DDA). A full scan was acquired at a target value of 1,000,000 ions with resolution R = 60,000 over mass range of 350-1600 amu. The top ten most intense ions were selected for fragmentation in the linear ion trap using Collision Induced Dissociation (CID) using a maximum injection time of 25 ms or a target value of 4000 ions.

The MS raw data (378 raw data files) were processed with MaxQuant v. 1.5.2.8^43^ and searched with Andromeda search engine^44^, querying two different UniProt databases^45^: Homo sapiens (09/07/2016; 92,939 entries) and Mus musculus (20/06/2016; 57,258 entries). Protein hits coming from individual database were separated using “Split protein groups by taxonomy ID” option in MaxQuant. The “Re-quantify” and “Match between runs” options were also enabled. For quantitation, multiplicity was set to 2 and Arg0 and Lys0, and Lys8 and Arg10 were used for ratio calculation of SILAC labelled peptides. Only unique peptides were used for protein group quantification. Digestion mode was set to “Specific” using the digestion enzyme trypsin and allowing maximum two miscleavages. Iodoacetamide derivative of cysteine was specified as a fixed modification, whereas oxidation of methionine and acetylation of proteins N-terminus were specified as variable modifications. First and main searches were carried out with precursor mass tolerances of 20 and 4.5 ppm respectively, and the MS/MS tolerance was set to 0.5 Da for the CID data. The FDR for peptide and protein identification was set to 1%; peptides with less than seven amino acid residues were excluded from processing. Only protein groups identified with at least one unique peptide were used for quantitation. The proteingroups.txt output file was analysed in Perseus^46^ v. 1.5.2.4. Reverse and contaminants hits (as defined in MaxQuant output), were removed from the list of identified proteins and only proteins quantified in at least 2 of 3 biological replicates in at least one tumour type were kept for further analysis. A median normalisation was performed on all samples before a Welch’s t-test with permutation based FDR set at 1% was used to identify significantly regulated proteins.

### Proteomic analysis of SLFN5-depleted tumours

Proteins obtained from total cell lysates were reduced with dithiothreitol and alkylated with iodoacetamide. Alkylated proteins were then precipitated in two steps using 24% and 10% solution of trichloroacetic acid. In both steps, pellets were incubated at 4°C for 10 minutes and centrifuged at 15.000 × g for 5 minutes. Supernatants were carefully aspirated and pellets washed with water until the supernatant reached neutral pH. Pellets were reconstituted and digested with endoproteinase Lys-C (Alpha Laboratories, UK) for 1 hour at room temperature and trypsin (Promega, Madison, WI, USA) overnight at 35°C.

Digested peptides were desalted using StageTip^47^ and separated by nanoscale C18 reverse-phase liquid chromatography performed on an EASY-nLC 1200 (Thermo Fisher Scientific, Waltham, MA, USA) coupled to an Orbitrap Fusion Lumos mass spectrometer (Thermo Fisher Scientific, Waltham, MA, USA). Elution was carried out using a binary gradient with buffer A (water) and B (80% acetonitrile), both containing 0.1% of formic acid. Peptide mixtures were separated at 300 nl/min flow, using a 50 cm fused silica emitter (New Objective, Woburn, MA, USA) packed in-house with ReproSil-Pur C18-AQ, 1.9 μm resin (Dr Maisch GmbH, Ammerbuch, Germany). Packed emitter was kept at 50°C by means of a column oven integrated into the nanoelectrospray ion source (Sonation, Germany). The gradient used start at 2% of buffer B, kept at same percentage for 3 minutes, then increased to 23% over 180 minutes and then to 32% over 40 minutes. Finally, a column wash was performed ramping to 95% of B in 10 minutes followed by a 5 minutes re-equilibration at 2% B for a total duration of 238 minutes. The eluting peptide solutions were electrosprayed into the mass spectrometer via a nanoelectrospray ion source (Sonation, Germany). An Active Background Ion Reduction Device (ABIRD, ESI source solutions, Woburn, MA, USA) was used to decrease ambient contaminant signal level. Samples were acquired on an Orbitrap Fusion Lumos mass spectrometer (Thermo Fisher Scientific, Waltham, MA, USA). The mass spectrometer was operated in positive ion mode and used in data-dependent acquisition mode (DDA). Advanced Peak Determination was turned on and Monoisotopic Precursor Selection was set to “Peptide” mode. A full scan was acquired at a target value of 4e5 ions with resolution R = 120,000 over mass range of 375-1500 amu. The top twenty most intense ions were selected using the quadrupole, fragmented in the ion routing multipole, and finally analysed in the Orbitrap, using a maximum injection time of 35 ms or a target value of 2e4 ions.

The MS .raw files were processed with MaxQuant software^43^ v. 1.6.3.3 and searched with Andromeda search engine^44^, querying SwissProt Homo sapiens database (30/04/2019; 42,438 entries). The database was searched requiring specificity for trypsin cleavage and allowing maximum two missed cleavages. Methionine oxidation and N-terminal acetylation were specified as variable modifications, and Cysteine carbamidomethylation as fixed modification. The FDR for peptide and protein identification was set to 1%.

MaxQuant proteingroups.txt output file was further processed using Perseus software v. 1.6.2.3^46^. The common reverse and contaminant hits (as defined in MaxQuant output) were removed. Only protein groups identified with at least one uniquely assigned peptide were used for the analysis. For label-free quantification, proteins quantified in all 3 replicates in at least one group were measured according to the label-free quantification algorithm available in MaxQuant^48^. Significantly regulated proteins were selected using Student’s t-test with a permutation based FDR of 5%.

### Transcriptomic analysis

Frozen tumours were manually crushed, reduced into powder and further processed using QIAshredder homogeniser columns (Qiagen, Hilden, Germany) before extraction. For cells, RNA was extracted 72 hours after initial seeding, when cells reached around 80% confluence. RNA extraction was performed using RNeasy Mini Kit (Qiagen, Hilden, Germany) with on-column DNase digestion (RNase-Free DNase Set, Qiagen, Hilden, Germany). Quality of the purified RNA was tested on an Agilent 2200 Tapestation using RNA screentape.

Libraries for cluster generation and DNA sequencing were prepared following an adapted method from Fisher et al^49^ using Illumina TruSeq Stranded mRNA LT Kit (Illumina, San Diego, CA, USA). Quality and quantity of the DNA libraries was assessed on a Agilent 2200 Tapestation (D1000 screentape) and Qubit (Thermo Fisher Scientific, Waltham, MA, USA) respectively. The libraries were run on the Illumina Next Seq 500 using the High Output 75 cycles kit (2 x 36 cycles, paired end reads, single index; Illumina, San Diego, CA, USA).

FastQ files were generated from the sequencer output using Illumina’s bcl2fastq (v. 2.20.0.422) and quality checks on the raw data were done using FastQC (v. 0.11.7) and FastQ Screen (v. 0.11.4)^50^. Alignment of the RNA-Seq paired-end reads was to the GRCh38^51^ version of the human genome and annotation using Tophat (v. 2.1.0)^52^. Expression levels were determined and statistically analysed by a workflow combining HTSeq (0.11.2)^53^, the R environment (v. 3.5.0, https://www.R-project.org), utilising packages from the Bioconductor data analysis suite^54^ and differential gene expression analysis based on the negative binomial distribution using the DESeq2 package^55^. Further data analysis and visualisation used R and Bioconductor packages.

### Metabolomic analysis

10^6^ cells were seeded in 6-well plates. The next day, medium was replaced and cells were allowed to grow for 48 hours. Cells were then washed 3 times with ice-cold PBS and metabolites were extracted by adding 1 ml of ice-cold extraction buffer (50% Methanol, 30% acetonitrile, 20% H_2_O). Plates were incubated on a shaker at 4°C for 5 minutes and supernatant was collected, centrifuged at 16,000 × g for 10 minutes and finally transferred to HPLC glass vials. Samples were kept at −80°C prior to LC-MS analysis.

The LC-MS method has been described previously^56^. Briefly, data were acquired using a Q Exactive Orbitrap mass spectrometer (Thermo Scientific, Waltham, MA, USA) coupled with a Thermo Ultimate 3000 HPLC system. The HPLC setup consisted of a ZIC-pHILIC column (SeQuant, 150 × 2.1mm, 5μm, Merck KGaA, Darmstadt, Germany), with a ZIC-pHILIC guard column (SeQuant, 20 × 2.1mm) and an initial mobile phase of 20% 20 mM ammonium carbonate, pH 9.2, and 80% acetonitrile. 5 μl of samples were injected and metabolites were separated over a 15 minutes mobile phase gradient, decreasing the acetonitrile content to 20%, at a flow rate of 200 μl/min and a column temperature of 45°C. All metabolites were detected across a mass range of 75-1000 m/z using the Q Exactive mass spectrometer at a resolution of 35,000 (at 200 m/z), with electrospray (ESI) ionization and polarity switching to enable both positive and negative ions to be determined in the same run. The mass accuracy obtained for all metabolites was below 5 ppm. Data were acquired with Thermo Xcalibur software.

The peak areas of different metabolites were determined using Thermo TraceFinder v. 4.0 software where metabolites were identified by the exact mass of the singly charged ion and by known retention time on the HPLC column. Commercial standards of all metabolites detected had been analysed previously on this LC-MS system. Final data were normalised to protein content, determined using the BCA protein assay kit (Thermo Fisher Scientific, Waltham, MA, USA).

### Immunohistochemistry and analysis

All patients involved in this study gave their written informed consent. These studies were conducted in accordance with recognized ethical guidelines UBC CREB number: H09-01628 and the amendment has been reviewed by the Chair of the University of British Columbia Clinical Research Ethics Board and the accompanying documentation was found to be acceptable on ethical grounds for research involving human subjects. Biochemical relapse was defined according to the ASTRO definition and represents three consecutive rise of detectable PSA following surgery.

To assess SLFN5 protein levels, immunohistochemistry was conducted with the Ventana DISCOVERY Ultra (Ventana Medical Systems, Tucson, Arizona, USA), an automated staining platform. Formalin-fixed paraffin-embedded (FFPE) TMA sections were baked, deparaffinized, and incubated in antigen retrieval solution CC1 (Ventana) at 95°C for 64 minutes. Following, anti-SLFN5 antibody (rabbit, 1:100, ab121537, abcam,) was incubated at room temperature for 1 hour. For detection, UltraMap DAB anti-Rb Detection Kit (Ventana) was used. Stained slides were scanned with Leica Aperio AT2 (Leica Microsystems, Concord, Ontario, Canada). The area of interest in the tumour images were delineated by pathologist. Positively stained cells were quantified with Aperio ImageScope (Leica Biosystems, Buffalo Grove, Illinois, USA).

### Proximity Ligation in situ Assay

22Rv1 cells were cultured on coverslips, washed with cold PBS and fixed with cold methanol for 20 minutes at −20°C. All incubations were performed in a humidity chamber. Blocking was done with 0.1% tween-TBS with 1% BSA for 1 hour at 37°C. Anti-ATF4 (#E4QAE, Cell Signaling) and SLNF5 (#121537, abcam) primary antibodies were incubated overnight at 4°C. P-LISA staining was performed as previously described^57^. Briefly, PLA Probe Anti-Mouse PLUS (DUO92001, Sigma-Aldrich, France) and PLA Probe Anti-Rabbit MINUS (DUO92005, Sigma-Aldrich, France) were used for secondary probe-linked antibodies. Ligation and Amplifications steps were performed according to manufacturer’s recommendations. All steps were interspersed by several washes using 0.1% tween-TBS. Nuclei were stained with DAPI (4’,6’-diamidino-2-phénylindole) and slides mounted with Fluoromount Aqueous Mounting Medium (F4680, Sigma Aldrich, France) before analysis using Zeiss LSM microscope confocal.

### Western Blot

Ten to thirty micrograms of proteins were loaded on to pre-casted SDS-PAGE gels (Invitrogen, Thermo Fisher Scientific, Waltham, MA, USA) and transferred to a PVDF membrane (GE Healthcare, Chicago, IL, USA). Following blocking, the membrane was probed overnight with primary antibodies (Supplementary Table 1) diluted in 5% milk-TBST. The next day the membrane was washed with TBST and incubated with respective fluorophore- or HRP-conjugated secondary antibodies diluted in 5% milk-TBST. For revelation, membrane was either scanned on a LI-COR Odyssey CLx Imaging system (LI-COR Biosciences, Lincoln, NE, USA) or revealed using Pierce™ ECL Plus Western Blotting Substrate (Thermo Fisher Scientific, Waltham, MA, USA) followed by image acquisition on a MyECL machine (Thermo Fisher Scientific, Waltham, MA, USA).

### qPCR analysis

RNA was extracted from cell or tumour samples using the RNeasy Mini Kit (Qiagen, Hilden, Germany) with on-column DNase digestion (RNase-Free DNase Set, Qiagen, Hilden, Germany). cDNA was prepared from 4 μg RNA using High-Capacity cDNA Reverse Transcription Kit (Thermo Fisher Scientific, Waltham, MA, USA). Real-time PCR was performed on the ABI 7500 FAST qPCR system (Thermo Fisher Scientific, Waltham, MA, USA) using TaqMan Universal Master Mix (Thermo Fisher Scientific, Waltham, MA, USA) and Universal ProbeLibrary probes (Roche, Basel, Switzerland). *CASC3* was used as a normaliser. Data are expressed as relative levels compared to control cells. The primers used in this study are listed in Supplementary Table 2.

### Immunofluorescence

50.000 cells were seeded on 19-mm coverslips and allowed to grow for 72 hours. Cells were then washed with ice-cold PBS and fixed with a 1:1 Acetone/Methanol solution for 20 minutes at −20°C. Cells were subsequently washed, incubated in blocking solution (10% FBS, 0.5% BSA, 0.3% Triton X-100 in PBS) and probed overnight with primary antibody solution (1% BSA, 0.3% Triton X-100, 2μg/mL SLFN5 antibody in PBS) at 4°C. The next day, secondary antibody solution was added (1% BSA, 0.3% Triton X-100, 1:100 anti-rabbit Alexa 555 antibody in PBS) for 1 hour at room temperature. Finally, slides were mounted with a DAPI-containing mounting solution (Thermo Fisher Scientific, Waltham, MA, USA). Pictures were acquired with a Nikon A1R Z6005 (Nikon Instruments, Melville, NY, USA) and processed using ImageJ (v. 1.46r).

### Analysis of SLFN5 binding motifs

Analysis of putative SLFN5-binding motifs^18^ in the promoter regions of *SLC7A5*, *SLC3A2*, *ATF4*, *NDNF* and *STRBP* was done with the Motif-based sequence analysis tools FIMO package^58^ (MEME Suite v. 5.1.1). Promoter regions were defined as 2000bp upstream of the transcription start site using the ensemble GRCh38.93 genome assembly.

### Statistical analysis

Statistical analyses were performed using GraphPad PRISM software v. 8.4.2 (GraphPad Software Inc, San Diego, CA, USA).

### Data reproducibility

Figure 1: Panel **D**: representative image from 3 independent biological experiments.

Figure 2: Panel **A:** n = 1 gel loaded with three tumour orthografts per condition. Panel **B:** representative image from 3 tumour orthografts per condition. Panels **C, E, F**: representative image from 3 independent biological experiments. Panel **D, G**: n = 6 (3 independent biological experiments performed in duplicates). Panel **I**: n = 3 independent biological experiments.

Figure 3: Panels **A, B**: n = 151; 162; 45; 29 for untreated; NHT-treated; CRPC and NEPC respectively. Panel **C**: n = 38 for high SLFN5 expression and n = 47 for low SLFN5 expression. Panel **D**: n = 70 for Gleason score < 7 and n = 56 for Gleason score >7. Panel **E**: n = 153 for non-metastatic patients and n = 75 for metastatic patients.

Figure 4: Panels **A**, **B**, **C**: representative image from 3 independent biological experiments. Panel **C**: n = 6 (3 independent fields taken from two independent migration inserts per condition). Panels **D**, **E**: representative image from 8, 9, 7 tumours for CTL, KO1 and KO2 respectively. Panels **F**, **G**: n = 8, 9, 7 tumours for CTL, KO1 and KO2 respectively.

Figure 5: Panel **A**: representative image from 3 independent biological experiments. Panel **D**: n = 6 (3 independent biological experiments performed in duplicates).

Figure 6: Panels **A**, **B**, **C**, **D**, **E**: representative image from 3 independent biological experiments. Panel **F**: n = 6 (3 independent biological experiments performed in duplicates). Panels **H**: representative image from 3 independent biological experiments.

Figure 7: Panel **A**: n = 3 independent biological experiments. Panels **B**, **C**: representative image from 3 independent biological experiments. Panel **D:** n = 1 gel loaded with four tumour orthografts per condition.

## Supporting information

Supplementary data 1_Proteomics CRPC

Supplementary data 2_RNAseq SLFN5 KO

Supplementary data 3_Pathway enrichment RNAseq

Supplementary data 4_SLFN5 binding sites

Supplementary data 5_ATF4 binding sites

Supplementary Table 1_Antibodies

Supplementary Table 2_Primers

Table 1_Significantly downreg genes

## Data availability

The raw files and the MaxQuant search results files have been deposited as partial submission to the ProteomeXchange Consortium via the PRIDE partner repository *[Perez-Riverol Y, Csordas A, Bai J, Bernal-Llinares M, Hewapathirana S, Kundu DJ, Inuganti A, Griss J, Mayer G, Eisenacher M, Pérez E, Uszkoreit J, Pfeuffer J, Sachsenberg T, Yilmaz S, Tiwary S, Cox J, Audain E, Walzer M, Jarnuczak AF, Ternent T, Brazma A, Vizcaíno JA (2019) The PRIDE database and related tools and resources in 2019: improving support for quantification data. Nucleic Acids Res 47(D1):D442-D450 (PubMed ID: 30395289)]* with the dataset identifiers PXD021405 and PXD021428. The following databases were used in this study: The Cancer Genome Atlas (TCGA—https://tcga-data.nci.nih.gov/tcga/); GEPIA (http://gepia.cancer-pku.cn/); STRING v11.0 (https://string-db.org/cgi/input.pl). All the data supporting the findings of this study are available within the article and its supplementary information files and from the corresponding author upon reasonable request.

## Acknowledgements

We would like to thank the Core Services and Advanced Technologies at the Cancer Research UK Beatson Institute, with particular thanks to the Metabolomics and Proteomics Units, and the PRIDE team. This work was supported by Cancer Research UK Beatson Institute core funding (C596/A17196) and CRUK core group awarded to HYL (A15151) and to SZ (A29800). This project has received funding from the European Union’s Horizon 2020 research and innovation programme under the Marie Skłodowska-Curie grant agreement No 721746. PP and EH were funded by grants from “La ligue Contre le Cancer”, “la région Bourgogne Franche-Comté” and “Canceropole Grand Est”. RSM is a Horizon ITN Early Stage Researcher. MS is a Medical Research Council Clinical Research Fellow (MR/L017997/1). CN is the recipient of CRUK Clinical Research Fellowship (grant 300444-01).

## Author contributions

RSM, MJS, AB and HYL designed the study. RSM, MJS, LR, GRB, WC, EH, ER, SHYK, LCAG, CNi, SL, DS, AB performed the experiments. RSM, MJS, CN, AH, PP, EH, SL, GMM, LF, DS, AB analysed the data. RSM, PP, DS, MEG, SZ, AB and HYL interpreted and discussed the data. AB and HYL wrote the manuscript. All authors critically reviewed the manuscript.

## Supplementary Figure Legends

**Supplementary Figure 1:**
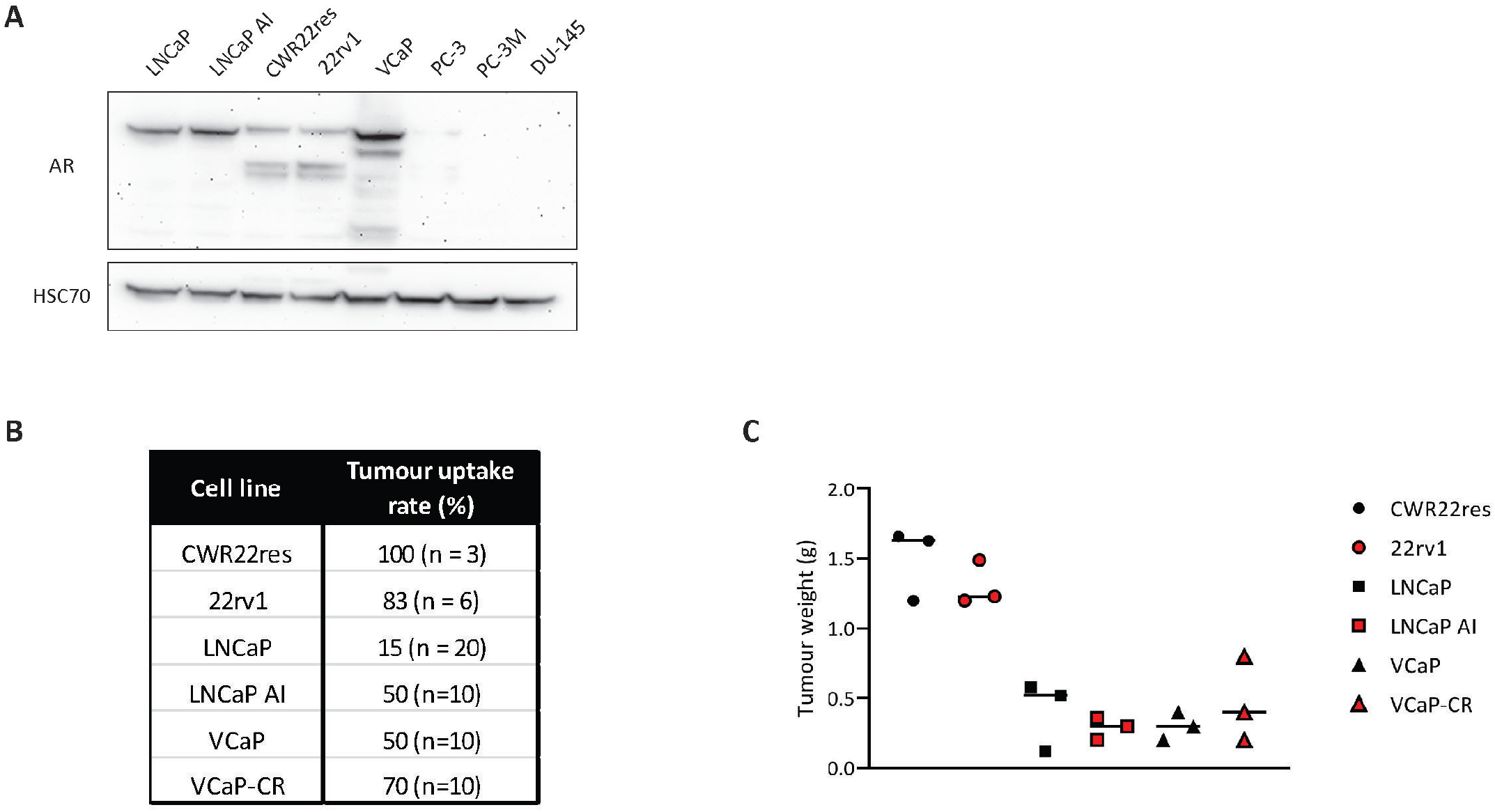
Characterisation of *in vivo* CRPC orthografts. **A**, Western blot analysis of AR expression in prostate cancer cell lines. HSC70 was used as a sample loading control. **B**, Percentage of in vivo tumour uptake from different prostate cancer cells following intra-prostatic injection. **C**, Tumour weight of CRPC orthografts in comparison to their respective HN counterparts. Panel **C**: Data are presented as mean values +/− SD. Panel **C**: n = three tumours per condition.

**Supplementary Figure 2:**
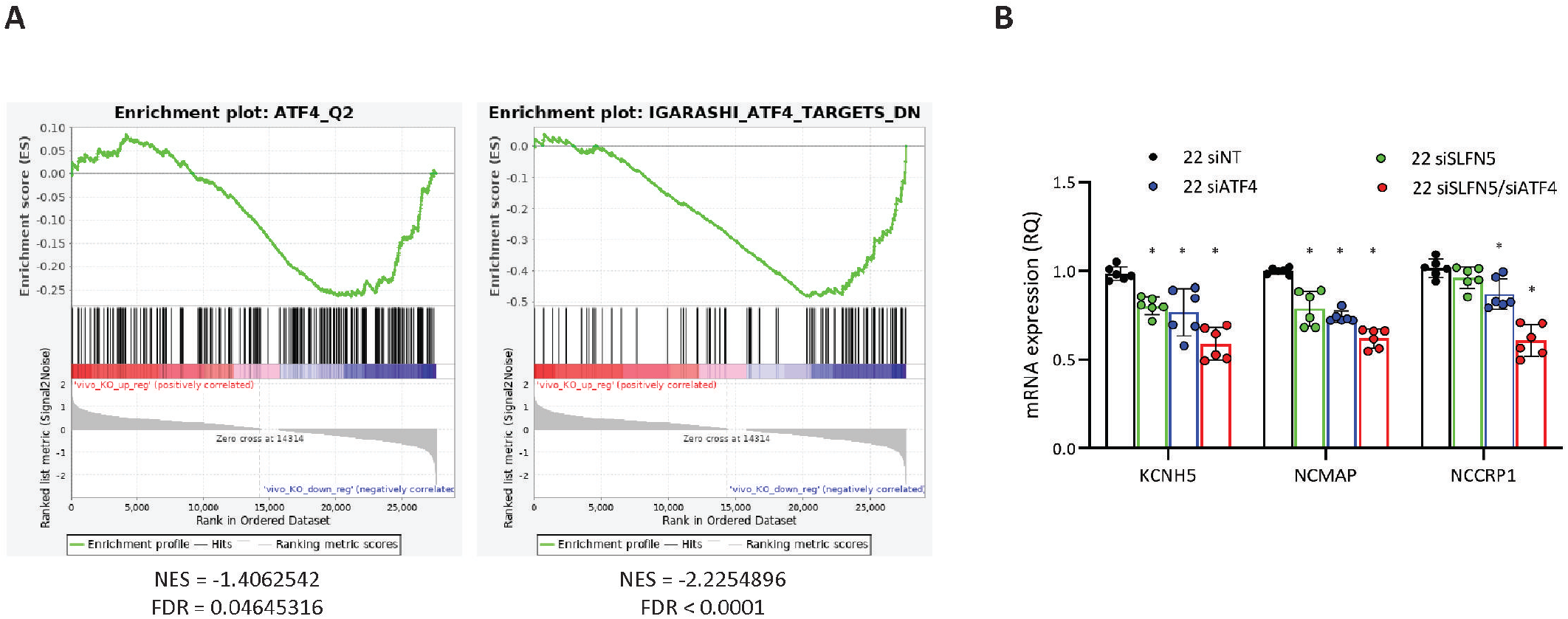
SLFN5 KO affects ATF4 signalling *in vivo*. **A**, Gene set enrichment plots for two additional ATF4-related gene sets analysed from SLFN5-depleted tumours transcriptomics. **B**, RT-qPCR analysis of *KCNH5*, *NCMAP* and *NCCRP1* expression in 22rv1 cells silenced for SLFN5, ATF4 or both. Panel **B**: Data are presented as mean values +/− SD. Panel **B**: *p-value < 0.05 using a 1-way ANOVA with a Tukey’s multiple comparisons test.

## Tables

**Table 1:** Significantly down-regulated genes (FC ≤ −3, p < 0.05) in SLFN5-depleted cells and tumour orthografts.

## Supplementary Datasets

**Supplementary Data 1:** Proteomic analysis of CRPC tumour orthografts.

**Supplementary Data 2:** RNAseq analysis of SLFN5-depleted cells and tumour orthografts.

**Supplementary Data 3:** Pathway enrichment analysis performed on RNAseq analysis of SLFN5-depleted cells.

**Supplementary Data 4:** Analysis of potential SLFN5-binding sites in the promoter regions of *SLC7A5*, *SLC3A2*, *NDNF*, *STRBP* and *ATF4*.

**Supplementary Data 5:** Analysis of potential ATF4-binding sites in the promoter regions of SLFN5-regulated genes (highlighted in Figure 5b).

