## Supplementary data 5_ATF4 binding sites for "Schlafen family member 5 (SLFN5) regulates LAT1-mediated mTOR activation in castration-resistant prostate cancer"

| **Gene symbol** | **Localisation**  **hg38** | **Sites**  **-5000 +1000** | **p-value** | **Reference** |
| --- | --- | --- | --- | --- |
| NDNF | chr4  121070535–121074535 | **-4108** | **0.0001** |  |
| TFPI | chr2  :187552435–187556435 | **-1241** | **0.0001** |  |
| KCNH5 | chr14  63043427–63047427 | **-2467** | **0.0001** |  |
| LUM | chr12  91109490–91113490 | **-2181** | **0.0001** |  |
| NCMAP | chr1  24554120–24558120 | +906  -3705  -4960 | 0.001  0.001  0.001 |  |
| SLC401A | chr2  189578783–189582783 | **-4427** | **0.0001** |  |
| CCDC68 | chr18  54957455–54961455 | +669,  -1074  -1199  -2129  -3233  -3928  -4240  -4476  -4690  -4828 | 0.001  0.001  0.001  0.001  0.001  0.001  0.001  0.001  0.001  0.001 |  |
| CFAP61 | chr20  20050540–20054540 | **-3843** | **0.0001** |  |
| DCN | chr12  91177479–91181479 | +985  -31,  -148,  -1899,  -2066,  -2987,  -3044,  -4012 | 0.001  0.001  0.001  0.001  0.001  0.001  0.001  0.001 |  |
| NCCRP1 | chr19  39194963–39198963 | **+810** | **0.0001** |  |
| PCDH15 | chr10  54799202–54803202 | **-1935** | **0.0001** |  |
| ORAI2 | chr7  102434201–102438201 | **623** | **0.0001** |  |
| SLC7A5 | chr16  87867505–87871505 | **683**  -**1616**  -**4871** | **0.0001**  **0.0001**  **0.0001** | Tameire et al.,2019  Han et al., 2013 |
| FLRT2 | chr14  85528143–85532143 | -**2228**  -**2659**  -**2952** | **0.0001**  **0.0001**  **0.0001** |  |
| STRBP | chr9 123266586–123270586 | **-4009** | **0.0001** |  |
| ACACB | chr12  109129360–109133360 | +669  +335  -54  -274  -1830  -3845  -4299  -4366 | 0.001  0.001  0.001  0.001  0.001  0.001  0.001  0.001 |  |
| BMPR1B | chr4  94755908–94759908 | -403  -766  -1786  -2980  -4196 | 0.001  0.001  0.001  0.001  0.001 |  |
| SLC3A2 | chr11  62878895–62882895 | -955  -1252  -2329  -2330 | 0.001  0.001  0.001  0.001 | Tameire et al.,2019 |
| FOXN4 | Chr12  109307242-109311242 | **-380** | **0.0001** |  |
| CRNKL1 | Chr20  20050466-20054466 | -3615  -1421  -1130  606 | 0.001  0.001  0.001  0.001 |  |

ATF4 couples MYC-dependent translational activity to bioenergetic demands during tumour progression. Tameire F, Verginadis II, Leli NM, Polte C, Conn CS, Ojha R, Salas Salinas C, Chinga F, Monroy AM, Fu W, Wang P, Kossenkov A, Ye J, Amaravadi RK, Ignatova Z, Fuchs SY, Diehl JA, Ruggero D, Koumenis C. Nat Cell Biol. 2019 Jul;21(7):889-899. doi: 10.1038/s41556-019-0347-9.

ER-stress-induced transcriptional regulation increases protein synthesis leading to cell death. Han J, Back SH, Hur J, Lin YH, Gildersleeve R, Shan J, Yuan CL, Krokowski D, Wang S, Hatzoglou M, Kilberg MS, Sartor MA, Kaufman RJ. Nat Cell Biol. 2013 May;15(5):481-90. doi: 10.1038/ncb2738.
