## Supplementary Table 1_Antibodies for "Schlafen family member 5 (SLFN5) regulates LAT1-mediated mTOR activation in castration-resistant prostate cancer"

| **Target** | **Species** | **Company** | **Reference** |
| --- | --- | --- | --- |
| 4EBP1 | Rabbit | Cell Signaling | 9644S |
| AR | Rabbit | Santa Cruz | sc-816 |
| HSC 70 | Mouse | Santa Cruz | sc-7298 |
| LC3 | Rabbit | Cell Signaling | 12741P |
| p-4EPB1 (Thr37/46) | Rabbit | Cell Signaling | 2855S |
| P70 | Rabbit | Cell Signaling | 9202 |
| p-P70-S6k (Thr389) | Rabbit | Cell Signaling | 9234S |
| p-S6 (Ser235/236) | Rabbit | Cell Signaling | 4856 |
| p-S6 (Ser240/244) | Rabbit | Cell Signaling | 2215 |
| SLC3a2 | Rabbit | Sigma | SAB1400263 |
| SLC7a5 | Rabbit | Cell Signaling | 5347S |
| SLFN5 | Rabbit | Abcam | ab121537 |
| Tubulin-α | Mouse | Santa Cruz | sc-8035 |

List of antibodies used in this study.
