## Supplementary Table 2_Primers for "Schlafen family member 5 (SLFN5) regulates LAT1-mediated mTOR activation in castration-resistant prostate cancer"

| **Gene** | **Fwd Sequence** | **Rev Sequence** | **Probe** |
| --- | --- | --- | --- |
| **CASC3** | ggggttccagttaatacaagtttc | gccagctgtatttctcttctgag | 84 |
| **SLFN5** | agcaagcctgtgtgcattc | ctggctggcagatgtttttc | 84 |
| **AR** | gccttgctctctagcctcaa | ggtcgtccacgtgtaagttg | 14 |
| **SLC7a5** | ttatacagcggcctctttgc | tgatcatttcctctgtgacga | 88 |
| **SLC3a2** | agccaaggctgacctcct | aggcgttccagctcaaga | 20 |
| **NCCRP1** | tgacgaacaaccagccatta | cagcagccagacatgcag | 42 |
| **NDNF** | gtcaaaacctgcagaaagca | catccagcaggtaagatttgc | 41 |
| **TFPI** | gcctgggcaatatgaacaat | ccacctggaaaccattcg | 47 |
| **KCNH5** | tttttggagaacatcgtcagg | caggccaatccacaatctg | 48 |
| **NCMAP** | gggggataccaccttcttct | tgatgatgacaaccaccaca | 50 |
| **STRBP** | aacaaggggagcttttgttg | ggggcagctgtgctgtaa | 68 |
| **ORAI2** | gatggaagtgcttggatgc | aggcacgttaagctcagcac | 58 |
| **ATF4** | tctccagcgacaaggctaa | ccaatctgtcccggagaa | 76 |

List of primers used in this study.
